# A scalable approach for genome-wide inference of ancestral recombination graphs

**DOI:** 10.1101/2024.08.31.610248

**Authors:** Árni Freyr Gunnarsson, Jiazheng Zhu, Brian C. Zhang, Zoi Tsangalidou, Alex Allmont, Pier Francesco Palamara

**Affiliations:** Centre for Human Genetics, University of Oxford, UK; Department of Statistics, University of Oxford, UK; deCODE genetics/Amgen, Reykjavík, Iceland; Doctoral Training Centre, University of Oxford, Oxford, UK

## Abstract

The ancestral recombination graph (ARG) is a graph-like structure that encodes a detailed genealogical history of a set of individuals along the genome. ARGs that are accurately reconstructed from genomic data have several downstream applications, but inference from data sets comprising millions of samples and variants remains computationally challenging. We introduce Threads, a threading-based method that significantly reduces the computational costs of ARG inference while retaining high accuracy. We apply Threads to infer the ARG of 487,409 genomes from the UK Biobank using ∼10 million high-quality imputed variants, reconstructing a detailed genealogical history of the samples while compressing the input genotype data. Additionally, we develop ARG-based imputation strategies that increase genotype imputation accuracy for ultra-rare variants (MAC ≤10) from UK Biobank exome sequencing data by 5-10%. We leverage ARGs inferred by Threads to detect associations with 52 quantitative traits in non-European UK Biobank samples, identifying 22.5% more signals than ARG-Needle. These analyses underscore the value of using computationally efficient genealogical modeling to improve and complement genotype imputation in large-scale genomic studies.

## Introduction

The ancestral recombination graph (ARG) is a graph in which nodes represent the genomes of a set of individuals or their ancestors, and edges represent genealogical connections between them. At any genomic position, the genealogical connections encoded in the ARG form a single genealogical tree, which changes along the genome due to recombination events that occurred in the transmission of genetic material from ancestors to descendants. The ARG efficiently integrates all these marginal trees into a single object and has been widely studied^1–3^. ARGs have also been leveraged in a wide range of genomic analyses, including generating synthetic data^4–7^, analyzing demographic history^8–11^ and natural selection^8,11^, detecting complex trait associations^12–14^, and facilitating polygenic prediction^15^.

Inference of ARGs from genotype data is computationally challenging due to the vast search space of graph topologies and coalescence times that could give rise to the observed genotypes^16^. For this reason, most methods rely on a combination of probabilistic inference and computational heuristics^8,10–13,17–20^. Recent approaches have improved the inference and analysis of genome-wide ARGs from sparse genotyping array data in large biobank data sets^13^. However, current methods struggle to scale to biobank-sized collections comprising millions of samples and genomic variants. Furthermore, recent work has shown that genealogical modeling may be used to improve genotype imputation accuracy^13,21^, but these potential benefits have not yet been attained in modern genomic data sets.

We introduce a scalable method for inferring ARGs, called Threads, which can be applied to biobank-scale datasets of genotyped, imputed, or sequenced individuals. We use extensive simulations to show that Threads requires significantly less computation and memory compared to other ARG inference methods and remains accurate despite relying on several modeling simplifications. We apply Threads to infer a genome-wide genealogy for 487,409 genomes from the UK Biobank using ∼10 million high-quality imputed variants. The resulting ARG integrates both genotypic and genealogical information about the analyzed samples, and an encoding derived from threading operations allows for more compact storage of input genotype data compared to commonly used genotype formats. Finally, we develop strategies that use genealogies inferred using Threads to increase genotype imputation accuracy of ultra-rare variants and use the inferred ARG to complement genotype imputation in association analyses of non-European UK Biobank samples.

## Results

### Overview of Threads

Threads infers ARGs through a process called *threading*^11,13^, whereby new haploid genomes are sequentially grafted onto a partial ARG by computing a set of “threading instructions”^13^. At each position along the genome, and for each sequence, these indicate a closest genetic relative (or cousin) from among samples already in the ARG, as well as a coalescence time. Once inferred for the whole sample, these threading instructions uniquely identify an ARG^13^. The threading instructions output by Threads also optionally specify whether the sequence and the closest genetic relative share the same allele. When allele sharing information is included, threading instructions are also sufficient to recover the input genotypes without assembling the ARG, as shown in Supplementary Figure 1.

Threads iteratively builds an ARG by considering all individuals in a fixed order and inferring the threading instruction for each of these individuals, as depicted in Figure 1. For each target individual, the threading instruction is computed with respect to the samples that have been added to the ARG in a previous iteration. To achieve high scalability, Threads breaks down the inference of threading instructions into three steps, which may be followed by an additional step that uses these instructions to assemble the ARG. First, Threads uses a haplotype matching algorithm based on the positional Burrows-Wheeler transform^22^ (PBWT) to select, for each haplotype, a set of candidate matches with an index lower than that of the target sample. Next, once all such candidate sets have been built, one among the set of most closely related haplotypes is selected at each site from among the candidate haplotypes by running the Li- Stephens algorithm^23^ in parallel across multiple samples. Finally, coalescence times are inferred using a likelihood-based approach based on the length of matching segments, the number of mismatching alleles, and the demographic history of the samples. A more detailed description of the Threads algorithm can be found in the Supplementary Note.

**Figure 1.**
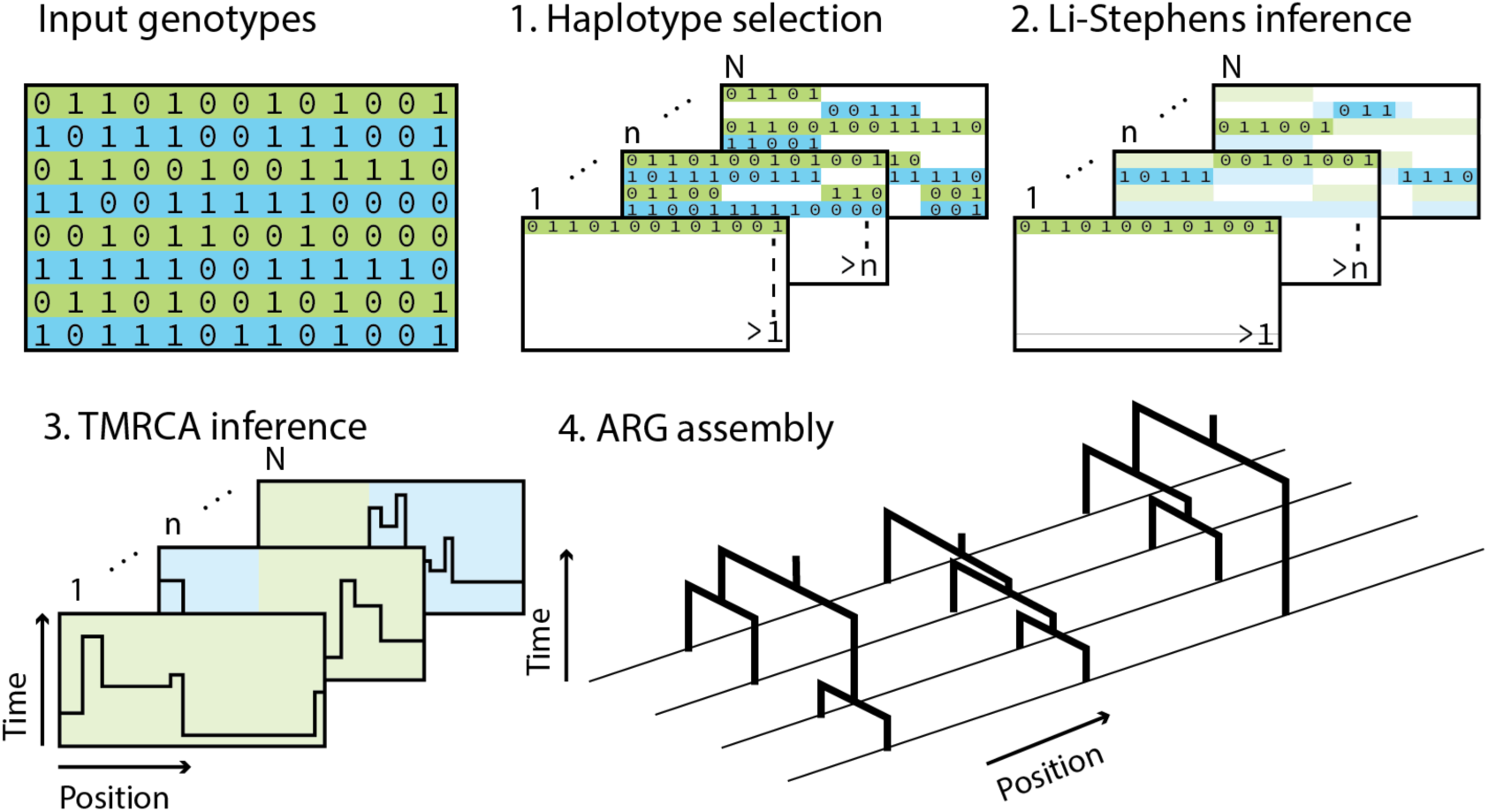
Overview of the Threads algorithm. Inference of threading instructions is performed in three steps. First, assuming a fixed order of input haplotypes, we select a subset of candidate genealogical closest cousins for each sample in each 0.5 centimorgan-sized window from among the samples of lower index. Second, we use the Li-Stephens algorithm to select a threading target (or genealogical closest cousin) from among the localized subsets. Third, we estimate the age of each segment inferred by the Li-Stephens algorithm. Finally, the inferred threading instructions may be used to assemble an ARG.

### Accuracy and scalability of Threads in simulations

We performed extensive simulations to test the computational scalability and accuracy of Threads. We included three other ARG inference methods in these benchmarks: Relate^8^, tsinfer^19^ combined with tsdate^10^ (tsinfer+tsdate), and ARG-Needle^13^, which is primarily optimized for inference from genotyping arrays and was only evaluated in that setting. We simulated sequencing and genotyping array data for up to 16,000 diploid genomes over a 15 megabase (Mb) region. The results of these analyses are summarized in Figure 2; additional results may be found in Supplementary Figures 2-10.

**Figure 2.**
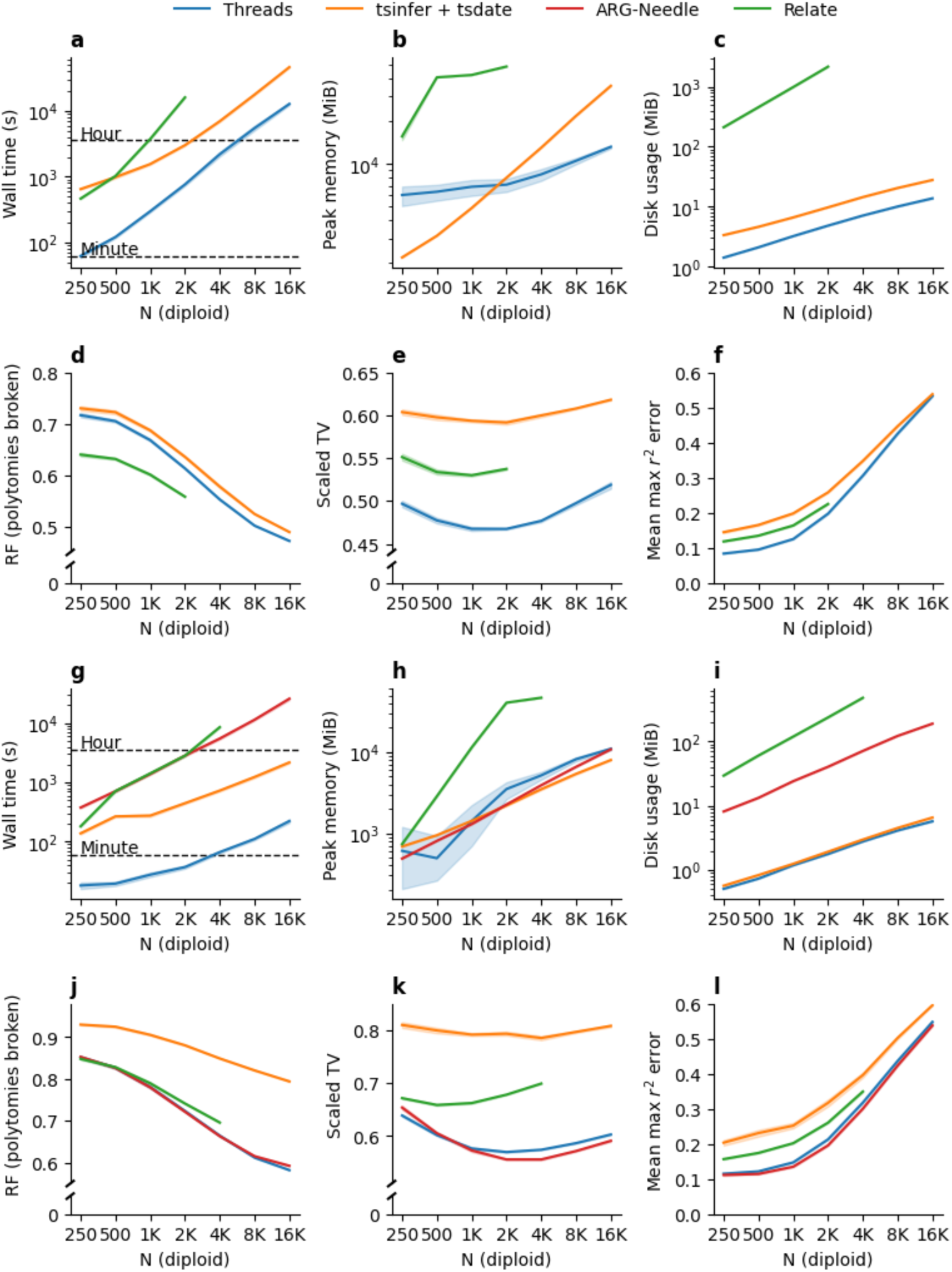
Computational cost and inference accuracy of Threads, tsinfer+tsdate, ARG- Needle, and Relate in simulated data. We simulated 15 Mb regions under a European demographic model with a constant recombination rate for sample sizes up to 16,000 diploid individuals. Evaluation of Relate was truncated at 2,000 samples for sequencing simulations and 4,000 samples for genotyping array simulations due to computational constraints. Shaded regions show bootstrap 95% confidence intervals based on 15 random seeds. **a-c.** Computational efficiency measured using runtime, memory consumption, and disk usage to store results, excluding temporary files written by Relate. **d-f.** The Robinson-Foulds (RF) metric, scaled total variation (TV), and mean max-r^2^ error compared with the ground truth ARG. Polytomies were randomly broken when evaluating the RF metric (see Methods). **g-i.** Runtime, memory consumption, and disk usage, excluding temporary files written by Relate, for the same simulated regions with markers subsampled to genotyping array density. **j-l.** RF, TV, and mean max-r^2^ error of ARGs inferred from simulated genotyping array data.

We first measured runtime, memory usage, and disk space used to store results in all simulation settings. All methods were provided with 8 computational cores for parallel computation. At its slowest, Threads ran ∼3 times faster than tsinfer+tsdate, the second fastest method. In many settings, Threads achieved speed-ups over other methods of an order of magnitude or more, consuming less memory and disk space (Figure 2, Supplementary Figures 2-4). We observed similar improvements when only one CPU was made available to all methods (Supplementary Figure 5).

We used several metrics to evaluate the accuracy of Threads and the other tested approaches. Due to computational costs, only three of these metrics were applicable to these large sample sizes, namely the Robinson-Foulds metric^24^ (RF), a tree-based measure of topological accuracy, the tree-total variation^13^ (TV), which probabilistically measures accuracy of both topology and branch lengths, and the max-r^2^ score, a stochastic accuracy score we devised to capture the extent to which variants implied by edges in the ARG tag other underlying genomic variants (see Methods). Because the RF metric is affected by the presence of polytomies (i.e., ARG nodes with more than two descendants) these were randomly resolved when present.

We performed several other secondary analyses using four additional metrics. We assessed the number of mutations that are needed to heuristically map a held-out genomic variant to an inferred marginal tree (mapping score, MS; see Methods). For completeness, we also employed three other metrics that measure the Euclidean distance between vectors summarizing marginal trees in various ways. These included the root-mean-square error^13^ (RMSE), which evaluates the accuracy of inferring pairwise genealogical distances, the topological Kendall-Colijn distance^25^ (KC), which measures distances to the root from internal nodes and was evaluated using randomly resolved polytomies, and the split-size vector metric^26^ (SV), which assesses clade size similarity. Due to their higher computational requirements, these vector-based metrics were only tested in smaller experiments. We note that the KC and the SV metric have been observed to be particularly sensitive to features such as tree shape and balance^13,26^, which may influence their interpretability in downstream analyses.

For sequencing data, Threads outperformed other methods under both the TV and max-r2 scores, while Relate achieved the highest accuracy under the RF and MS metrics (Figure 2, Supplementary Figures 6, 10). In simulated genotyping array data, either Threads or ARG-Needle was the most accurate or tied for highest accuracy across all simulation settings (Figure 2, Supplementary Figures 8, 10). Threads proved robust to genotyping errors (Supplementary Figure 7) and to an artificially low mutation rate (Supplementary Figure 9). In smaller experiments involving vector-based metrics, Relate outperformed Threads and tsinfer+tsdate under RMSE and SV, whereas tsinfer+tsdate achieved the highest accuracy under KC.

Genealogical encodings of genotype data have been shown to enable compressing genotype information in simulated data^7^, although real data sets yielded lower compression rates^19^. The threading instructions output by Threads provide an alternative efficient encoding of both genealogical and genotypic data. In addition to uniquely determining the inferred ARG^13^, these instructions can be used to reconstruct the genotypic data used to infer it, as described in Supplementary Figure 1. We assessed disk space requirements for storing genotype data using compressed threading instructions from Threads, PLINK2^27^ *pgen* format, and compressed ARGs from tsinfer+tsdate in *tszip* format. We found Threads to provide efficient storage compared to these formats (Supplementary Figure 11), with variation across simulation settings and dataset size. Compressed threading instructions could be used to efficiently recover the input genotype data directly, without the need to convert into other ARG formats (Supplementary Figures 1, 12, 13), although working with an assembled ARG often led to faster data retrieval, depending on factors such as the genotyping error rate.

Taken together, these results demonstrate that Threads can infer large-scale genealogies using fewer computational resources than other ARG inference methods, while remaining highly accurate and producing a compact encoding of both genealogical and genotype data.

### Genome-wide genealogies for the UK Biobank and 1000 Genomes Project data sets

We applied Threads to infer genome-wide ARGs using genotyped, imputed, and sequenced genomic variants for up to 487,409 individuals from the UK Biobank (UKB) and for 2,261 individuals from the 1000 Genomes Project (1KGP). Consistent with simulations, we observed that the compressed threading instructions output by Threads efficiently stored both genealogical and genotypic information for both data sets (Table 1). Threads’ output was particularly space-efficient for an ARG inferred from 9,992,478 imputed variants for the UKB data set, requiring only 7.1% of the space used to store the input *pgen* file. For an ARG inferred in a subset of 711,755 SNP array variants and 337,464 unrelated white British samples, threading instructions required 38.9% of the space required by the *pgen* input file. When applied to the 1KGP data set using 1,227,802 sequenced variants from chromosome 20, Threads produced threading instructions that required 30% of the disk space needed for the input *pgen* file. This also corresponded to 30.4% of the space to used store an ARG inferred using tsinfer+tsdate and compressed using *tszip*, which took 10.8× more time to compute. By converting these threading instructions into ARGs and analyzing average coalescence times and genealogical relationships between different geographic groups, we found that these inferred genealogies effectively retained fine-grained ancestry information, recovering patterns of population structure across the UK (Supplementary Figure 14, Methods).

**Table 1.** Computational resources for the 1KG and UKB data sets. We report the computing time and disk usage required to infer and store genealogical encodings, which include genealogical and genomic data, as well as the disk usage for storing the input genomic data. We present results for (a) Threads (.threads compressed threading instructions), tsinfer (.tsz ARG format), vcf.gz, and pgen formats for 1,227,802 variants from Chromosome 20 for 2,261 individuals from the 1,000 Genomes Project; (b) Threads (.threads compressed threading instructions) and pgen formats for the UK Biobank data set, for 487,409 samples and 9,992,478 HRC-imputed genome-wide variants, as well as for a subset of 337,464 unrelated white British samples and 711,754 SNP genome-wide array variants.

| Algorithm/encoding | 1,000 Genomes Project<br>N=2,261; M=1,227,802 (Chr 20) |  |
| --- | --- | --- |
|  | Size (Mb) | Inference time (h)* |
| Threads (.threads) | 46.7 | 1.24 |
| tsinfer (.tsz) | 153.7 | 13.4 |
| vcf.gz | 155.5 | - |
| pgen | 78.6 | - |

**Table 1. Computational resources for the 1KG and UKB data sets.**
| Algorithm/encoding | HRC-imputed<br>N=487,409; M=9,992,478 |  | SNP array<br>N=337,464; M=711,754 |  |
| --- | --- | --- | --- | --- |
|  | Size (Gb) | Inference time (h)** | Size (Gb) | Inference time (h)* |
| Threads (.threads) | 40.3 | 4,449.1 | 12.1 | 192.2 |
| pgen | 563.5 | - | 31.1 | - |

\* 8 cpu threads
\*\* 16 cpu threads

### ARG-based ultra-rare variant imputation

Genealogical relationships between samples can be utilized for genotype imputation^28–31^, where genomic variants not directly observed in a target individual are inferred using close genetic relatives found in a sequenced reference panel. Current genotype imputation strategies build on the Li-and-Stephens^23^ approach, which is highly effective but relies on approximate genealogical modeling. In particular, this approach does not allow modeling scenarios where the age of variants being imputed is less than the time to the most recent common ancestor between reference and target individuals (Figure 3a), which may lead to a systematic overestimation of rare-variant genotype dosages^13,21^. We verified this phenomenon using simulations, where we observed inflated dosages for variants with a minor allele count (MAC) of up to about 10, approximately independent of panel size (Supplementary Figure 15).

**Figure 3.**
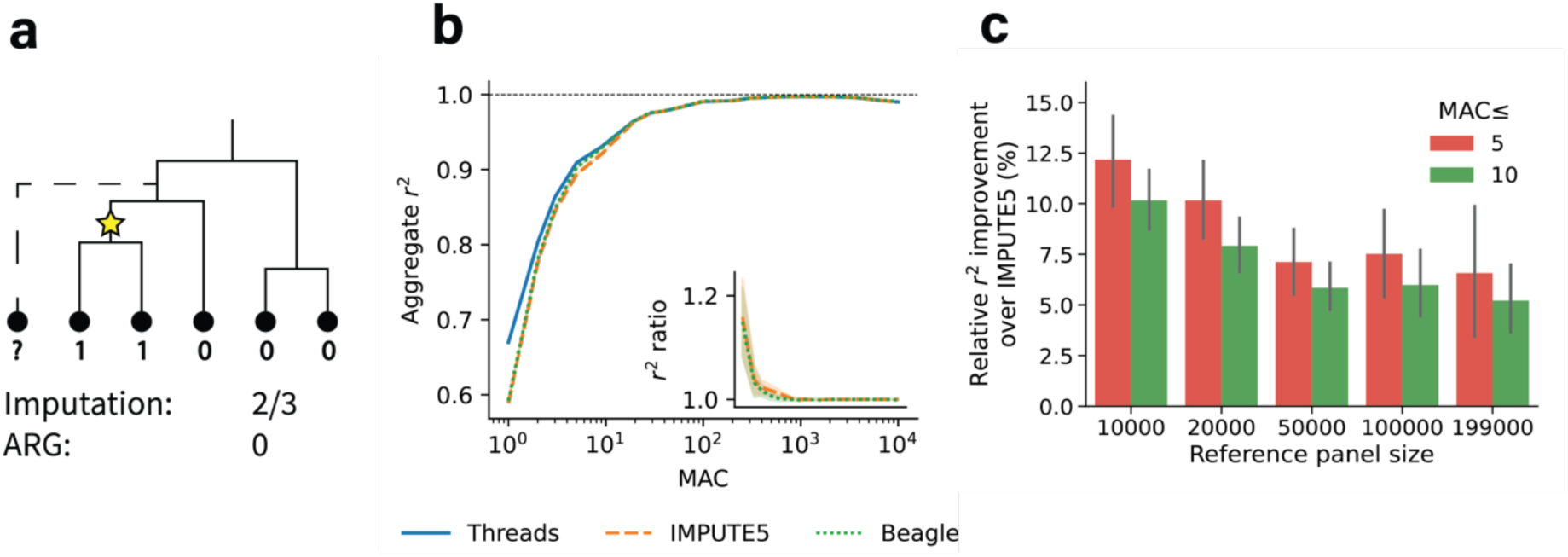
ARG-based genotype imputation. **a.** Toy example of how genotype dosages may be inflated even when the set of genealogical closest cousins is correctly inferred. Here, the target sample coalesces above the mutation (star symbol) carried by two of its closest cousins. **b.** Imputation quality for three methods using a simulated reference panel of 10,000 individuals. The inset shows the ratio between Threads and other methods across the same allele count spectrum. Shaded regions represent 95% confidence intervals across 10 random seeds. **c.** Relative improvement (%) in imputation accuracy of variants with MAC up to 5 or 10. We compare the results of applying Threads and IMPUTE5 to UK Biobank exome sequencing data. Error bars represent 95% confidence intervals computed using 260 regions of 10 cM each.

We developed an ARG-based imputation strategy that improves accuracy for ultra-rare variants when used with ARGs inferred using Threads. Briefly, we first inferred the ARG of a sequenced reference panel and assigned one or more associated edges in the ARG to variants in the reference panel. We then inferred probabilistic threading instructions for the target genome relative to the reference panel, estimating the probability that the target genome inherited each variant in the panel (Methods). This approach allows for modelling the possibility that the ancestor shared by the target and reference genomes is older than the age of the mutation being imputed. Since we expected imputation approaches based on the Li-Stephens algorithm to only result in inflated dosages for the rarest variants (Figure 3, Supplementary Figure 15), we adopted the Li-Stephens forward-backward algorithm^28,31^ for variants with a minor allele frequency exceeding 0.1% or minor allele count exceeding 20.

We assessed the accuracy of this threading-based genotype imputation approach through simulated and real datasets. We first simulated reference panels of up to 30,000 diploids and measured aggregate r^2^ for variants categorized by frequency in the panel. Threads showed a 10-15% improvement in imputation r^2^ for singleton variants compared to IMPUTE5 and Beagle 5.4. As allele counts increased, differences in accuracy decreased, with all methods reaching similar levels for variants of MAC=10 in the reference (Figure 3, Supplementary Figure 16). We next applied this method to an inferred ARG for up to 199,000 UKB exome-sequenced samples, obtaining accuracy improvements of ∼5-10% for variants with MAC between 1 and 10 (Figure 3). Finally, we applied this approach to African and European ancestry samples from the 1KGP data set, observing gains starting at 5-6% for the rarest variants, with the accuracy of all methods evening out at around MAC=10 (Supplementary Figure 17).

### ARG-based association testing

Recent work has shown that genealogy-wide association analyses, where genomic variants are inferred from a reconstructed ARG and tested for association against a heritable trait, can complement genotype imputation in the study of genomic variation that is not well represented in sequenced reference panels^13^. Inferred ARGs can also be used within a linear mixed model framework to estimate heritability^13^ by constructing an ARG-based genetic relatedness matrix (ARG-GRM) that may better capture unobserved genomic variation. When these ARG-GRMs are built for a specific genomic region, such as a gene, this approach can be used to perform ARG-based variance component association testing^14,32^.

We performed ARG-based association testing within the UK Biobank dataset, comparing the use of ARGs inferred using either Threads or ARG-Needle (see Methods). We focused on individuals of non-European genetic ancestry, who are underrepresented in imputation reference panels^33^, analyzing 8,235 unrelated individuals of Central/South Asian ancestry (CSA) and 6,253 unrelated individuals of African ancestry (AFR), as defined in the Pan-UKB project^34^. We tested for association between 21,378 protein-coding genes and non-coding RNA regions (Supplementary Table 1) with 52 blood cell indices and blood biochemistry marker levels (Supplementary Table 2), comparing several association strategies (see Methods). We performed variance component testing using an ARG inferred from ∼10 million HRC-imputed variants (Threads-HRC). Since ARG-Needle cannot be easily applied to data sets of this scale, we tested an ARG previously inferred using 711,754 SNP array markers^13^ (ARG-Needle-SNP). We also included tests performed using an ARG inferred using Threads for the same subset of markers (Threads-SNP). Finally, we compared these ARG-based approaches to standard association testing based on genotype imputation. To this end, we applied the same variance component association test that we used in ARG-RHE to test HRC-imputed variants (HRC-RHE), and ran standard mixed-model association testing of individual imputed variants using Regenie^35^.

In these analyses, ARG-based association effectively complemented imputation-based approaches (Figure 4; Supplementary Tables 3, 4; Supplementary Figure 18). Variance component association testing performed using ARGs inferred with Threads detected more gene-trait associations than when using ARGs inferred with ARG-Needle (N_Threads-HRC_ = 212, N_Threads-SNP_ = 182, N_ARG-Needle-SNP_ = 173, Figure 4), and more than when we applied the same variance component association test to imputed genotype data alone (N_HRC-RHE_ = 155). In addition, the signals detected using the ARG were complementary to those detected using imputation (Figure 4b,c, combined with HRC-RHE: N_Threads-HRC_ = 211, N_Threads-SNP_ = 208, N_ARG- Needle-SNP_ = 198, see Methods). This complementarity was also observed when comparing these associations to those detected using single-variant testing from a mixed-model analysis, with Threads again detecting a larger fraction of associations shared with imputation-based testing compared to ARG-Needle.

**Figure 4.**
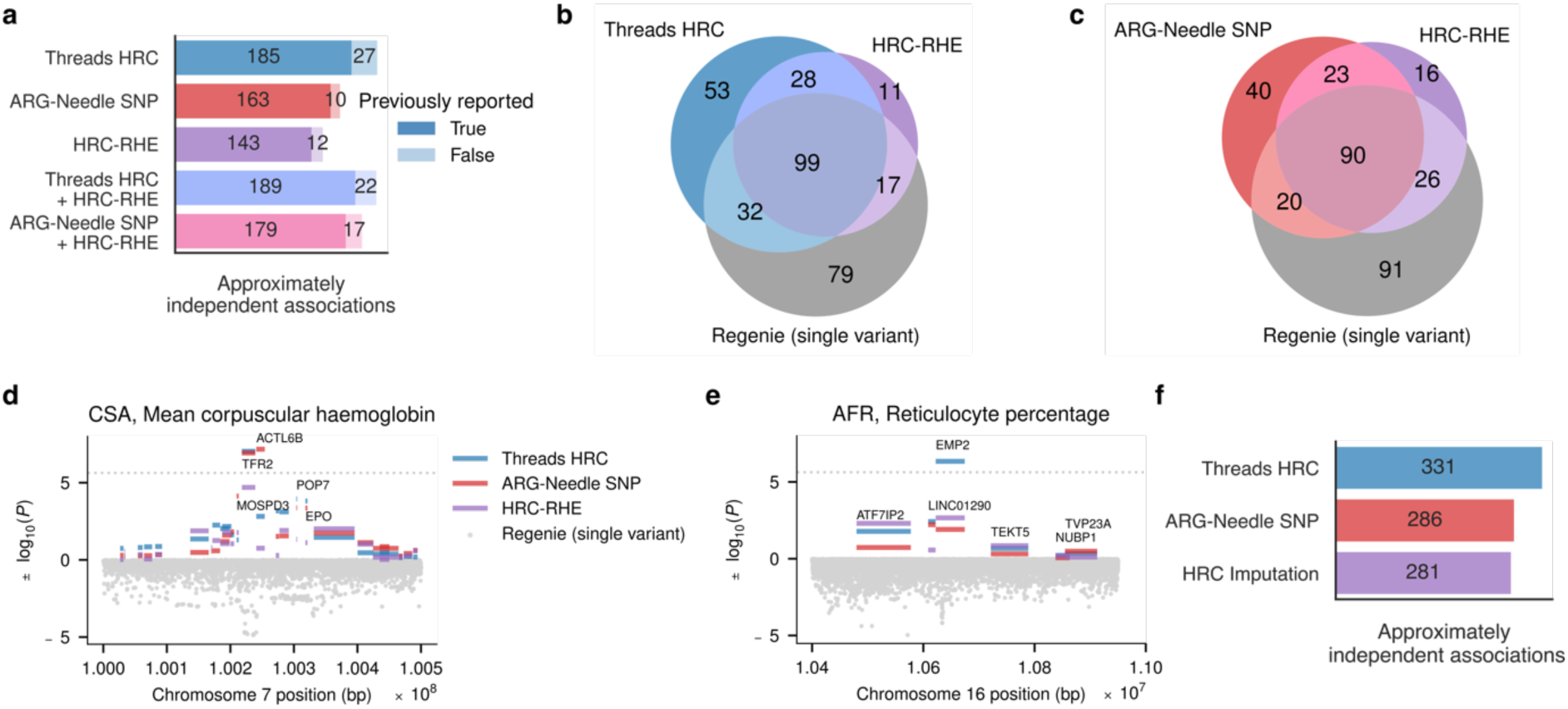
ARG-based association testing. **a.** Total numbers of LD blocks containing genome- wide significant gene-trait associations detected in the CSA and AFR subgroups using different methods. Each bar is annotated with the subsets of LD blocks containing gene-trait associations that were or were not previously reported on the Open Targets platform (see URLs). **b, c.** Overlap in LD blocks containing gene-trait associations detected using HRC-RHE, Regenie, and ARG-RHE applied to ARGs inferred using either Threads (**b**) or ARG-Needle (**c**). Combined counts from the CSA and AFR subgroups; group-specific counts are reported in Supplementary Figure 18. **d, e.** Manhattan plots for genomic regions containing the TFR2 and ACTL6B genes (**d**) in the CSA subpopulation for mean corpuscular hemoglobin level and the EMP2 gene (**e**) in the AFR subgroup for reticulocyte percentage. **f.** Approximately independent single-variant associations detected through genealogy-wide testing of ARGs inferred using Threads or ARG-Needle, as well as through genome-wide testing of HRC-imputed variants.

We verified these associations by checking for overlap with larger, more powered association studies, in which we found most signals detected in the CSA and AFR subgroups to be reported. Previously established signals which we detected using the ARG but not using HRC-imputed data alone in our analyses included, in CSA, associations between the TFR2, ACTL6B, and HFE genes and Mean Corpuscular Hemoglobin^36–41^, as well as between the SLC12A3 gene and HDL Cholesterol^39,42–44^. We also detected associations that did not overlap with those reported in the Open Targets database (see URLs), particularly for the AFR subgroup, for which fewer ancestry-matched large-scale studies exist. Among these, we detected associations between reticulocyte measures and EMP2, which has previously been associated to red blood cell counts and other blood traits^36,39,40,45^, and between MMP26 and neutrophil percentage, previously associated to lymphocyte counts, monocyte counts, and other blood traits^36,37,45–47^.

Overall, ARG-based association testing effectively complemented imputation-based approaches in these analyses, with Threads outperforming ARG-Needle in both the number of detected associations and their overlap with imputation-based signals. Although Threads is more computationally efficient, its reliance on model simplifications may reduce the accuracy of ARGs inferred from SNP data alone at deeper time scales. Consistent with this, when we used the ARG-MLMA approach to perform genealogy-wide mixed-model association testing of individual ARG-derived variants^13^, ARGs inferred by Threads using SNP data yielded fewer approximately independent associations than those inferred by ARG-Needle (see Methods, Supplementary Figure 18). Given that single-variant testing is less powered to detect associations with rare variants compared to variance-components-based testing^48^, this discrepancy likely reflects ARG-Needle’s higher accuracy in inferring common variants, which tend to originate from deeper time scales. However, Threads’ scalability allows it to be applied to denser collections of imputed variants, potentially improving the inferred ARG, particularly at deeper time scales. We note that genotype imputation may also introduce errors, which may affect the detection of long shared haplotypes. The effects of imputation on the accuracy of inferred ARGs may therefore vary across analyzed groups, reflecting variation in imputation accuracy for variants of different frequencies and ages.

## Discussion

We developed Threads, a scalable algorithm for the inference of ancestral recombination graphs. Through extensive simulation and benchmarking, we verified that Threads performs ARG inference using fewer computational resources than other available methods, while retaining high accuracy. We applied Threads to infer genome-wide ARGs for 2,261 samples from the 1,000 Genomes Project, using ∼57 million variants, and for 487,409 samples from the UK Biobank, using ∼10 million high-quality imputed variants. The inferred ARGs, encoded using compressed threading instructions, compactly store both genotype and genealogical data while requiring significantly less disk space compared to standard genotype formats. We developed strategies to use these inferred ARGs to perform genotype imputation, observing accuracy improvements for imputed ultra-rare variants (MAC ≤ 10) in both simulated and UK Biobank exome sequencing data. Finally, we used the ARGs inferred in non-European samples to detect associations in 52 complex traits, where Threads increased the number of associated loci compared to ARG-Needle and detected signals that complement those obtained using standard imputation-based strategies.

Our work highlights connections and potential synergies between genealogical inference and genotype imputation. The initial step of the Threads algorithm uses the PBWT data structure to rapidly identify genetic relatives who share long-range haplotypes, a strategy also used in genotype imputation algorithms^28^. Additionally, to compute the threading instructions used to reconstruct the ARG, Threads estimates coalescence times between samples. Our analyses demonstrate that this step can also be leveraged in genotype imputation to improve the accuracy of the rarest imputed variants. Similar benefits are likely to be observed in the closely related problem of haplotype phasing, where haplotype-based modeling of coalescence times has been shown to improve performance^49^. In a preliminary analysis, we used a simple algorithm that leverages inferred ARGs to phase ultra-rare variants. This approach was as accurate as SHAPEIT5 when applied to 1KGP data (Supplementary Figure 19b) but less accurate in other simulated scenarios (Supplementary Figure 19c,d), suggesting the need for further methodological development.

In addition to improving the imputation accuracy of rare variants, using an ARG to store the reference panel could facilitate the sharing of these panels. As recently suggested^50^, truncating an inferred ARG to remove genealogical connections within the most recent generations may preserve sufficient information to effectively perform analyses such as phasing and imputation, while offering some protection for the privacy of the individuals in the panel. This may facilitate data sharing, and the threading of new individuals onto these references could provide an effective algorithmic strategy towards building a more comprehensive global genealogical resource.

We identify several limitations and potential areas for future improvements. First, Threads estimates the age of genomic regions using a Li-Stephens algorithm, assuming these regions correspond to pairwise identical-by-descent (IBD) segments. While this approximation has proven reasonably accurate, particularly in large samples (see Supplementary Note), additional modeling may lead to improved accuracy for the ages inferred during the initial iterations of the algorithm, which typically influence the length of ARG edges in the deeper past. Furthermore, threading instructions inferred for the subset of samples that are initially threaded into an ARG can be easily replaced with another set of instructions for the same individuals. An inferred ARG may therefore be improved at a later stage by substituting these initial instructions with those derived from an ARG constructed using algorithms that are slower but more accurate in small data sets. Second, like other threading approaches, Threads incrementally adds new individuals to an existing ARG, resulting in slight variations in the ARG depending on the order of the individuals, which may be leveraged to obtain simple estimates of uncertainty^13^. The use of subtree pruning and regrafting operations^11^ may lead to improved uncertainty quantification. Third, although the threading instructions inferred by Threads allow for efficient compression of genotypes, many ARG-based operations can be inefficient when performed directly on these instructions. In this case, it may be beneficial to convert the threading instructions into other formats or assemble them into ARG data structures, which requires additional computational resources. Fourth, although we have shown that modeling of coalescence times can improve the imputation of ultra-rare variants, the current ARG-based imputation algorithm within the Threads package is at least an order of magnitude slower than standard, optimized algorithms. This underscores the need for further computational optimization of this approach. Fifth, Threads can more easily scale to analyses comprising large collections of imputed variants, but the quality of the inferred ARGs will depend on imputation accuracy, which in turn depends on several population- and sample-specific features. Despite these limitations and areas of future improvement, Threads provides a valuable new tool for the inference and analysis of genome-wide genealogies at biobank scales.

## Methods

For a detailed description of the Threads algorithm, please refer to the Supplementary Note.

### Simulations

To evaluate the accuracy of ARG inference methods, we simulated ARGs using msprime^51^ under two demographic models, a model inferred using SMC++^52,53^ for Northern Europeans from Utah (CEU) from the 1000 Genomes Project^54^ and a constant demographic model of effective population size *N_e_* = 10,000 diploid individuals. All simulations used a fixed recombination rate of 1.3 × 10^−8^per site per generation, approximately matching the average genome-wide recombination rate. We simulated mutations both using a realistic rate of 1.4 × 10^−8^, compatible with recent estimates^55^, and an artificially low mutation rate of 1.4 × 10^−9^. In each case we simulated sequences of 15 Mb in length with sample sizes ranging from 250 to 16,000 diploid individuals. To avoid boundary effects, all accuracy metrics were evaluated only on the central 5 Mb region. Each experiment was repeated for 15 different random seeds. Genotyping array data sets were simulated by subsampling the observed genotypes to match in frequency with genotyping array data from the UK Biobank. In experiments involving genotype imputation, we simulated regions of 20 Mb in length and evaluated accuracy on the central 10 Mb for 10 random seeds, but otherwise kept simulation parameters unchanged^54^.

### ARG metrics and ARG benchmarks

We evaluated seven tree- or ARG-based accuracy metrics for Threads, ARG-Needle^13^ (v1.0.1), Relate^8^ (v1.2.1), tsinfer^19^ (v0.3.1) and tsdate^10^ (v0.1.5). Evaluation of Relate was truncated at 2,000 samples for sequencing experiments and at 4,000 samples for genotyping array experiments due to computational constraints. We evaluated the Robinson-Foulds metric^24^ (RF), the ARG total variation^13^ (TV) and the max-r^2^ score for all sample sizes. In addition, we computed the topological Kendall-Colijn metric^25^ (KC), the root-mean-square error metric^13^ (RMSE), and the split-size vector metric^26^ (SV) for sample sizes up to 4,000. The mapping score was evaluated only at sample size 2,000, the largest sequencing simulations where all methods were included.

The max-r^2^ score is an ARG-based accuracy metric measuring the correlation between variants in an inferred ARG and underlying true variants, which reflects the extent to which an inferred ARG is expected to tag underlying variation in a genealogy-wide association study. Given a ground-truth ARG A and an inferred ARG B, we simulate 𝑀*_A_* and 𝑀*_B_* mutations on A and B respectively, by uniformly distributing them along the ARG volume. Then, for each mutation 𝑚*_A_* on A, represented as a bit-set, we compute max*_m_B__*_∈*B*_ 𝑟^2(^(𝑚*_A_*, 𝑚*_B_*), thus finding the mutation on B that is most closely correlated with 𝑚*_A_* from among the 𝑀*_B_* mutations on B. The max-r^2^ score is defined as the mean over all such correlations,

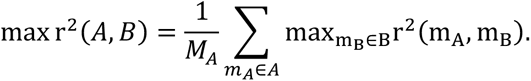

For the experiments of Figure 2 and Supplementary Figures 6-9, we set 𝑀*_A_* = 𝑀*_B_* = 10,000, so that the computational time required to evaluate this metric remained independent of sample size. Note, however, that this may cause the max-r^2^ score to increase with sample size, due to a corresponding increase in ARG volume. An alternative approach, which would be less computationally efficient but would allow keeping the sampling rate constant over ARG edges, is to set the number of resampled mutations to be proportional to ARG volume.

We define the mapping score (MS) of an observed variant, given an inferred marginal tree, as the minimum number of edges in the tree where mutations need to be added to cover all and only the carriers of the variant, divided by the total number of carriers. We compute the minimum number of edges using the following iterative approach. First, we select any variant carrier for which an ancestral mutation has not yet been found. We then consider the sequence of edges connecting this node to the root and add a mutation to the last edge in this sequence that only subtends individuals who carry the variant. We consider these subtended carriers to be covered by the mutation and iterate until all remaining carriers have been covered. We then compute the MS as the ratio between the number of added mutations and the number of carriers. In simulations, when a ground-truth ARG is known, we compute the mapping score by randomly generating new mutations under the ground-truth ARG and aggregating their mapping score by ancestral allele frequency.

### ARG inference in the UK Biobank and 1000 Genomes data sets

To infer genome-wide genealogies for the UK Biobank from markers imputed using the Haplotype Reference Consortium (HRC) reference panel^30,56^, we first extracted bi-allelic SNPs with INFO score ≥ 0.95 and minor allele frequency (MAF) ≥ 0.0001. We rounded all dosages within 0.1 of the nearest integer, otherwise setting genotypes to missing, and discarded all variants with missingness greater than 10%. All filtering was performed using PLINK2^27^. We divided the genome into chunks of 15 cM along with 1 cM of padding on each end and phased using SHAPEIT5-common^49^. We then joined pairs of chunks using SHAPEIT5-ligate, to create chunks of 30 cM, with 1 cM of padding. This procedure gave a total of 136 chunks, which were used to infer ARGs using Threads using parameters --query_interval 0.02 and -- match_group_interval 1.0 instead of the default 0.01 and 0.5, respectively. These parameters determine the density of queries for candidate matches in the haplotype matching step of the algorithm and may be raised to reduce memory usage; in our experiments Threads proved robust to the choice of matching algorithm parameters. To infer ARGs for samples of African and Central/South-Asian ancestry, we applied the same procedure, with a minor allele count (MAC) threshold of 1 and without any chunking, inferring ARGs for whole chromosome arms at a time using Threads with default parameters. For ARGs based on genotyping arrays, we followed the procedure described by Zhang et al.^13^ and phased the arrays using Beagle 5.1^57^ and divided the genome up into 166 chunks of equal size, inferring ARGs using Threads in genotyping array mode. To infer ARGs for 2,261 individuals from the 1,000 Genomes Project, we downloaded phased, curated data comprising 56,935,222 variants (see URLs) and applied Threads directly to each chromosome arm without any further chunking.

### ARG-based analysis of population structure

To extract regional structure (Supplementary Figure 14), we subsampled the ARG to 100 samples from each of 122 UK postcodes, keeping only self-identified white British individuals^56^. For each sample, we then counted the occurrences of each postcode within the set of genealogical closest cousins genome-wide, querying marginal trees of the ARG at 10- kilobase intervals. We averaged these observations across individuals within each postcode. Regional heatmaps show the proportion of closest genetic relatives from each postcode for a single, focal postcode. To visualize regional structure^19,58^, we focused on the ARG of 49,354 self-reported white British individuals^56^ from five postcodes in north-east England (DH, Durham; DL, Darlington; NE, Newcastle; SR, Sunderland; TS, Middlesborough) and evaluated the proportion of genome shared between individuals within 10 generations. We averaged these observations across individuals and divided by the number of observations to obtain an affinity matrix that was used for average-linkage hierarchical agglomerative clustering algorithm as implemented in scikit-learn^59^ (v.1.3.0). We truncated the clustering operation at different relatedness thresholds, ranging from 𝛼 = 0.1 down to 𝛼 = 1 × 10^−5^, extracting the largest clusters for each 𝛼.

### Threading-based compression

For experiments quantifying the disk space required by Threads’ output, we measured the disk space required to store compressed threading instructions. These threading instructions consist of a partition of a genomic region into discrete segments, and attached to each such segment, a threading target, a coalescence time, and a list of sites on the segment where the sequence and the threading target are heterozygous. These quantities are serialized and then compressed using the HDF5 format^60^. In addition to uniquely determining the ARG^13^, these threading instructions can be used to recover the input genotype variants, as they define, at each site, a directed acyclic graph that can be used to extract genotypes, as illustrated in Supplementary Figure 1.

### Threading-based imputation

We evaluated the accuracy of threading-based genotype imputation in both simulations and real data. We simulated reference panels and genotyping array data sets of size ranging from 100 to 30,000 samples for 10 random seeds using the parameters described above and used these to impute 10 held-out target samples. We estimated accuracy using the aggregate r^2^ binned by minor allele count in the panel. In experiments involving the 1,000 Genomes Project data, we used 10 held-out samples of European ancestry (2 from each of 5 sub-populations: CEU, Northern Europeans from Utah; FIN, Finnish; GBR, British; IBS, Iberian; TSI Tuscans from Italy) and 7 held-out samples of African ancestry (1 from each of 7 sub-populations: ACG, African Caribbean in Barbados; ASW, African-American in Soth West USA; ESN, Esan in Nigeria; YRI, Yoruba, Nigeria; LWK, Luhya, Kenya; GWD, Gambian; MSL, Mende, Sierra Leone) from a set of 2,261 unrelated samples from the 1,000 Genomes Project^54,61^. In experiments using the UK Biobank exome sequencing data, we evaluated imputation accuracy on 100 held-out samples using a reference panel of up to 199,000 exome sequenced samples.

To perform threading-based imputation, we first inferred an ARG for the reference panel using Threads. We then mapped variants to edges in the ARG using a method previously described by Speidel et al.^8^. This approach can be applied in cases where the ancestral allele status is unknown and accounts for errors in the input genotypes and the inferred ARG by allowing for imperfect overlap between the leaves subtended by lineages assigned to mutations and their observed carriers. This enabled us to estimate the age *s* of a mutation as the midpoint of its mapped lineage. As we only applied ARG-based imputation to rare variants, we heuristically assumed the major allele to be ancestral. For variants that did not map to the ARG, we performed standard imputation using the Li-Stephens model. Next, we inferred threading instructions with respect to the reference ARG for each sample being imputed. We modeled the age of a shared haplotype, *t*, as an Erlang-2 distribution with parameter 𝜆, a function of segment length and population size (see Supplementary Note). We used this to compute the threading dosage:

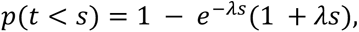

which estimates the probability that the lineage being threaded inherited the mutation. To account for uncertainty in the threading target, we used the PBWT to recover all sequences that are identical to the threading target along the segment being imputed. Finally, to account for uncertainty on the boundaries of the segment, and to further account for uncertainty in the threading target, we also ran a forward-backward algorithm to get a full posterior matrix *P*, with 𝑃*ij* denoting the probability that sample *i* is the genealogical closest cousin for the target at site *j*. To obtain the dosage for each site, we computed the expectation

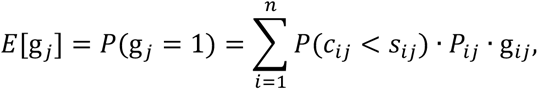

where *n* is the number of sequences in the reference panel. Here, 𝑃(𝑐*_ij_* < 𝑠*_ij_*) denotes the probability that at site *j* the target sequence coalesces to reference sample *i* at time 𝑐*_ij_* lower than the age 𝑠*_ij_* of the mutation carried by sample *i* at site *j*, with 𝑠*_ij_* = 0 if it carries no mutation. The term g*_ij_* denotes the observed genotype of sample *i* at site *j* in the panel.

### ARG-based phasing

To phase rare variants using the ARG, we traversed a marginal coalescence tree upwards, starting from each of the two possible haplotypes carrying an unphased variant, until at least one haplotype of all heterozygous carriers and both haplotypes of all homozygotes were found. We defined the average tree-based distance to all such carriers as the *phasing distance* for the haplotype and placed the variant on the haplotype of lower phasing distance. More formally, if for each unphased carrier *c*, *c*_1_ and *c*_2_ are its two haplotypes, then the phasing distance can be written as

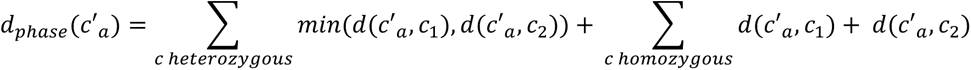

for *a* = 1,2. If *c*’ is the only carrier of the mutation, we set 𝑑*_phase_*(𝑐′*_a_*) to be the negative of the age of its most recent common ancestor. The singleton case prioritizes the haplotype that is less related to the rest of the sample and more likely to carry a singleton mutation. A similar step based on identical-by-state tracts is performed by SHAPEIT5^49^. For all ARG-based analyses, we rely on common-variant scaffolds inferred using SHAPEIT5 for variants with MAF above 0.1%.

### Association analyses

We tested 52 quantitative blood traits including 27 blood cell indices and 25 blood biochemistry marker levels (see Supplementary Table 1). We applied standard preprocessing steps^45^ to these phenotypes separately for the AFR and CSA subgroups. We first stratified samples based on sex and menopause status and applied a rank-inverse-normal transformation (RINT). We then regressed out covariates, which included alcohol use, smoking status, age, age squared, height, body mass index (BMI), assessment center, genotyping array, and 20 genetic principal components, and applied RINT a second time. We considered 21,378 protein-coding and non-coding RNA regions spanning more than 1,000 bp from the University of California, Santa Cruz (UCSC) genome browser Gene Nomenclature Committee (HGNC) table (see Supplementary Table 2 and URLs).

We performed gene-based, variance component association testing using the ARG-RHE algorithm^32^ implemented in the arg-needle-lib library (v1.0.3). We modeled the vector of phenotypes 𝑦 for 𝑛 individuals as 𝑦 ∼ 𝑁(0, 𝜎*_g_*^2^𝐾 + 𝜎*_g_*^2^𝐼), where 𝐾 is a GRM formed by variants within a region, 𝐼 is an 𝑛 × 𝑛 identity matrix, and 𝜎*_g_*^2^ + 𝜎*_e_*^2^ = 1 are scalars reflecting genetic and environmental variance components. In the case of ARG-based association testing, the GRM 𝐾 is a local Monte Carlo ARG-GRM from normalized resampled mutations within a region, built as in as in Zhang et al.^13^ by sampling variants with MAC ≥ 5 from the ARG at a rate 𝜇 = 10^−6^. For HRC-RHE, 𝐾 is the local GRM using normalized imputed genotypes in a gene region having MAC ≥ 5, INFO score > 0.3, missingness < 0.1, and Hardy-Weinberg equilibrium p > 10^-15^. We used the ARG-RHE algorithm to perform variance component association testing, using GRMs built from ARG-derived (referred to as ARG-RHE) or imputed (referred to as HRC-RHE) variants within each of the considered gene regions (Supplementary Table 1). We applied a Bonferroni correction at a 5% family-wise error rate, obtaining a genome-wide significance threshold of 0.05/21,378 ≈ 2.3 × 10^-6^ for each trait. We also report the results of combining the ARG-RHE and HRC-RHE tests on both ARG-derived and imputed variants, by comparing the smaller p-value for the gene given by the two tests with a genome-wide significance threshold of 0.05/21,378/2 ≈ 1.2 × 10^-6^.

As a baseline, we also performed single-variant association testing using Regenie^35^, testing HRC-imputed dosages on the same 52 preprocessed phenotypes with covariates regressed out. We applied the same variant filtering criteria as in the HRC-RHE analysis, which led to ∼30 million imputed variants, and adopted a genome-wide significance threshold obtained through resampling-based testing^13,62^ (see Supplementary Table 5). To count associated regions for each approach, we assigned genome-wide significant signals into approximately independent LD blocks provided by LDetect^63^, which include 2,583 and 1,443 LD blocks for the AFR and CSA subgroups respectively.

We also performed genome-wide and genealogy-wide association analyses for the same preprocessed traits, following the methodology of Zhang et al.^13^. For GWAS, we kept variants with MAC ≥ 5, missingness < 10%, and INFO score > 0.3 and then performed association using BOLT-LMM (v.2.3.4), without excluding related individuals. For genealogy-wide association, we inferred ARGs from SNP array and imputed data as described above. We then tested for association using the same parameters as Zhang et al.^13^, including a sampling rate of 𝜇 = 10^−5^ and calibration factors estimated by BOLT-LMM, while filtering out clades with derived allele count < 5. Genome-wide significance thresholds for this analysis, reported in Supplementary Table 5, were established using resampling-based testing^13,62^. To estimate the number of approximately independent associations, we first performed stringent clumping using PLINK^27^ (v.1.90) with r^2^=0.01. We then ran a conditional-joint analysis on the clumped SNPs using GCTA-COJO^64,65^ with default parameters, setting the --cojo-slct flag with --cojo-p 1e-7.

## Data availability statement

UK Biobank data can be accessed by approved researchers through https://www.ukbiobank.ac.uk/. The 1000 Genomes Project data set can be accessed through http://ftp.1000genomes.ebi.ac.uk/vol1/ftp/data_collections/1000G_2504_high_coverage/working/20220422_3202_phased_SNV_INDEL_SV/. Variant annotations were obtained through the genome aggregation data base (https://gnomad.broadinstitute.org/), coding and non-coding RNA regions were obtained through the UCSC genome browser (https://genome.ucsc.edu/cgi-bin/hgTables), known variant associations were obtained through the Open Targets GWAS association database (https://genetics.opentargets.org), ancestry allocations for the UK Biobank were downloaded from the Pan-UKB project (https://pan.ukbb.broadinstitute.org/).

## Code availability statement

The Threads software is available at https://palamaralab.github.io/software/threads. External software can be downloaded from the following URLs: msprime (v.1.2.0 https://pypi.org/project/msprime/), tsinfer (v.0.3.1, https://pypi.org/project/tsinfer/), tsdate (v.0.1.5, https://pypi.org/project/tsdate/), Relate (v.1.2.1, https://myersgroup.github.io/relate/), ARG-Needle (v1.0.2, https://pypi.org/project/arg-needle/), IMPUTE5 (v.1.1.5, https://jmarchini.org/software/#impute-5), Beagle 5.4 (v.22Jul22.46e, https://faculty.washington.edu/browning/beagle/beagle.html), SHAPEIT5 (v.5.1.1, https://odelaneau.github.io/shapeit5/), GCTA (v.1.94.1, https://yanglab.westlake.edu.cn/software/gcta/), PLINK2 (v1.90b6.26 and v2.00a3.7LM, https://www.cog-genomics.org/plink/2.0/).

## Acknowledgements

We thank R. Fournier for discussions and suggestions. This work was conducted using the UK Biobank resources (application no. 43206) and supported by the Clarendon Scholarship (Á.F.G., B.C.Z.); the Keble College de Breyne Clarendon Scholarship (Á.F.G.); Wellcome Trust ISSF grant no. 204826/Z/16/Z (P.F.P); Wellcome Trust grant no. 222336/Z/21/Z (Á.F.G.); ERC Starting Grant ARGPHENO no. 850869 (P.F.P, B.C.Z.); EPSRC grant EP/S023151/1 (Z.T.); EPSRC Centre for Doctoral Training in Health Data Science (EP/S02428X/1) (J.Z.). Computation used the Oxford Biomedical Research Computing (BMRC) facility, a joint development between the Centre for Human Genetics and the Big Data Institute supported by Health Data Research UK and the NIHR Oxford Biomedical Research Centre. Financial support was provided by the Wellcome Trust Core Award Grant Number 203141/Z/16/Z. The views expressed are those of the author(s) and not necessarily those of the NHS, the NIHR or the Department of Health.

## Contributions

Á.F.G. and P.F.P. designed the Threads algorithm. Á.F.G implemented algorithms and performed simulations and analyses of 1000 Genomes Project data. Á.F.G, J.Z., and Z.T. performed analyses of UK Biobank data. B.C.Z and A. A. provided software tools. Á.F.G. and P.F.P. wrote the manuscript.

