## Supplementary Note for "A scalable approach for genome-wide inference of ancestral recombination graphs"

for Gunnarsson et al. "A scalable approach for

#### Contents

**1 Introduction** **2**

**2 Haplotype matching with the PBWT** **2**

**3 Memory-efficient Viterbi inference using the Li-Stephens model** **4**

**4 Dating path segments** **12**

**7 References** **26**

### 1 Introduction

This supplementary note provides additional details for the Threads algorithm, including inference heuristics and a discussion on simplifying assumptions. As described in the main text, Threads takes as input a set of phased genotypes and outputs as set of "threading instructions" for each sample. More precisely, for a genomic region $[L_1, L_2]$ and a set of genotypes indexed $1, \ldots, N$, Threads infers for each $n \leq N$ a map $[L_1, L_2] \to \mathbb{R}_+ \times \{1, \ldots, n-1\}$, representing, for each site, the coalescence time of sample $n$ and a threading target from among the previously threaded samples.

The threading instructions are obtained in two steps, first inferring the closest genealogical relative at each site and then the coalescence time. In what follows, we will refer to the closest genealogical relative both as the "closest cousin" and the "threading target". Closest cousin inference is itself split into two sub-routines, a haplotype matching algorithm based on the positional Burrows-Wheeler transform (PBWT) [7] followed by a memory-efficient and perfectly parallelizable implementation of the Viterbi algorithm under the Li-Stephens model [15]. Coalescence times are estimated using segment-wise likelihood-based modeling.

### 2 Haplotype matching with the PBWT

The Li-Stephens model can be used for closest cousin inference but a direct application to large-scale ARG inference by threading quickly becomes computationally unfeasible. As the sample size grows, each sample $n$ must search through $n-1$ previously threaded sequences for close matches, resulting in a time complexity of $O(MN^2)$ for the whole ARG inference process, where $M$ is the number of sites and $N$ the total number of samples.

To reduce the search space for the Li-Stephens algorithm, Threads first constructs a PBWT of the input genotypes [7] to find a small set of candidate matches for each sample being threaded. This strategy is related to haplotype matching methods employed by modern phasing and imputation algorithms to reduce the state space of a Li-Stephens model [3, 4, 11, 24], although there are several key differences compared to these approaches, which are detailed below.

Threads starts by partitioning the input region into small chunks to estimate candidate haplotype matches independently within each chunk. This allows the algorithm to maintain a locally optimized Li-Stephens state space which is important for inference over large genomic regions. The default genomic chunk size for this step is 0.5cM; we denote the number of obtained genomic chunks by $M_{\text{chunk}}$.

The haplotype matching then proceeds by sorting the input haplotypes at each site into a PBWT prefix array $a$. At regular intervals, the prefix array is queried for the $L$ neighbours around each sequence $n$ from among the sequences $1, \ldots, n-1$, keeping track of how many times each sequence has been chosen as a candidate match. More precisely, if sequence $n$ has prefix index $i$ at site $j$, i.e. $a_{i,j} = n$, then the first $L/2$ sequences $a_{k,j}$ immediately above and below $a_{i,j}$ in the prefix array satisfying $a_{k,j} < n$ are considered as candidate matches. If fewer than $L/2$ such indices $k > i$ or $k < i$ are found then the search window is expanded in the opposite direction until $L$ matches have been found or until all sequences have been considered. The default interval between these queries is $0.01$cM and the default window size is $L = 4$. We denote by $M_{\text{query}}$ the number of sites queried for haplotype matches. Threads does not keep track of the PBWT divergence array. Instead, it heuristically assumes neighbouring sequences in the PBWT prefix array share long tracts that are identical by state. A similar heuristic is employed by IMPUTE5 [24] for efficient genotype imputation.

The querying operation is implemented by sequentially inserting the prefix-array index of each sequence $n = 1, \ldots, N$ at the query site into a red-black tree, as implemented by the C++ container `std::set`, and querying the $L$-sized neighbourhood around the insertion index in the tree. The time complexity for the complete matching step thus becomes $O(MN + M_{\text{query}} N \log N)$. Pseudocode is provided in Algorithm 1. In practice, we do not include singleton variants in this step of the analysis as they tend to have higher phasing and genotyping errors than other variants [3, 11] and may interfere with identification of long stretches of identical-by-state regions.

Finally, once all sites within a chunk have been processed, the candidate sequences are filtered by the number of times they were matched within the chunk.

By default, we discard any matches observed in fewer than 4 queries. If no matches remain after this filter, the threshold is decreased by 1 until at least one sequence is matched. For $n < 100$, we include all sequences observed in the querying step and for $n \geq 10,000$, we double the number of matches required for inclusion, to heuristically limit the number of candidates and save on memory usage. Moreover, we include the top 4 matched sequences from adjacent chunks, to account for the possibility that the chunk breakpoint lies close to a recombination event.

The full PBWT or the full genotype matrix are never stored in memory. Instead, genotypes are streamed through the algorithm in small chunks, only keeping track of the match candidates for each sequence. For each sample, the maximum possible number of matches found per chunk is $LM_{\text{query}}/M_{\text{chunks}}$, although in practice it is usually much lower. Thus, the complete haplotype matching step can be performed using only a single pass through the data, with a total memory footprint of $O(NLM_{\text{query}})$.

### 3 Memory-efficient Viterbi inference using the Li-Stephens model

In this section we present the memory-efficient Viterbi algorithm used in Threads for the inference of threading targets. For a reference set $\mathcal{H}$ of $N$ haplotypes over $M$ sites and a query haplotype $g$ over the same sites, the Li-Stephens model [15] uses prior information on mutation and recombination rates in the reference sample to model $g$ as a mosaic of haplotypes from $\mathcal{H}$, minimizing the number of allelic mismatches and recombinations. This is achieved by assigning a probability $P(\pi)$ to each path $\pi \in \{1, \ldots, N\}^M$ through the reference panel. A path through $\mathcal{H}$ of maximum probability can be found in $O(NM)$ time and memory using the Viterbi algorithm and we call such a path a *Viterbi path*.

In classical implementations of the Viterbi algorithm, a full per-site, per-sample matrix of probabilities is constructed before a traceback step is performed to extract a best path. This approach is computationally prohibitive in modern biobank data sets, which can contain hundreds of thousands of samples and millions of

**Algorithm 1** Threads haplotype matching for a single genotype chunk

**Input:** A genotype matrix $X \in \{0,1\}^{M \times N}$ and query sites $Q$.

**Output:** For each $n = 1, \ldots, N$, haplotype matches from $\{1, \ldots, n-1\}$.

```
C ⇐ [{ }, ..., { }]                              ▷ Empty multi-set for each n = 0, ..., N − 1
sort_0 ⇐ [0, ..., N − 1]
sort_1 ⇐ [0, ..., N − 1]
for i = 0, ..., M − 1 do
    ac ⇐ ∑_j X[i, j]
    c_0, c_1 ⇐ 0
    for j = 0, ..., N − 1 do                     ▷ Compute the next PBWT sorting
        if X[i, j] = 1 then
            sort_1[N − ac + c_1] ⇐ sort_0[i]
            c_1++
        else
            sort_1[c_0] ⇐ sort_0[i]
            c_0++
        end if
    end for
    sort_0 ⇐ sort_1
    if i ∈ Q then                                ▷ Query site for matches
        p ⇐ [ ]                                  ▷ p[j] is the index of sample j in sort_0
        for j = 0, ..., N − 1 do
            p[sort_0[j]] ⇐ j
        end for
        Initialize an empty red-black tree t
        t.insert(p[0])
        for j = 1, ..., N − 1 do
            k ⇐ t.insert(p[j])                   ▷ k is the insertion index of p[j] in t
            matches ⇐ L-sized neighborhood around k in t
            for l ∈ matches do
                C[j].insert(s_0[l])
            end for
        end for
    end if
end for
return C
```

sites. The number of haplotypes under consideration can be reduced significantly by applying steps such as the haplotype matching described above, but even then the memory requirements for long genomic tracts are high. Threads relies on an algorithm that builds a Viterbi path using $O(NM)$ runtime and, on average, $O(N)$ memory. Here, we focus only on the memory-efficient Viterbi algorithm.

The memory-efficient Viterbi algorithm used within Threads (hereafter referred to as Threads-Viterbi) relies on a branch-and-bound approach based on two key observations. First, recombination events in the output of the Viterbi are rare, especially for large $N$, resulting in paths that consist of few but long segments. In Viterbi paths obtained using a reference panel of 2,251 unrelated sequences from the 1000 Genomes Project [1, 5, 9], we find that the average segment length is over 200 kilobases, and in genotyping array data from unrelated white British samples in the UK Biobank ($N$=337,464), the length of an average segment is well over a megabase. This observation allows us to store whole Viterbi paths using a small amount of memory, by only noting the segment breakpoints and the associated threading target. Second, under the Li-Stephens model, recombination events are symmetric, that is,

$$p(\pi_{i+1} = \beta | \pi_i = \alpha, \text{ recombination between } i \text{ and } i+1 \text{ }) = \frac{1}{N},$$

for any states $\alpha, \beta$. This property allows substantially reducing the search space for possible Viterbi paths, as more formally stated in the following Proposition.

**Proposition 1.** *Suppose $\pi^{(i)}$ is a Viterbi path for the subset of the panel $\mathcal{H}$ containing sites $1$ through $i$. If there exists a Viterbi path through $\mathcal{H}$ that recombines between sites $i$ and $i+1$, then there exists a Viterbi path $\pi$ through $\mathcal{H}$ satisfying $\pi_i = \pi_i^{(i)}$.*

More succinctly, at site $i$ we only need to consider recombination events from a sequence of highest probability given all observations up to $i$.

We now describe the Threads-Viterbi algorithm in detail. The algorithm consists of alternating "branch" and "bound" steps, with a single "traceback" step at the end. Branch-and-bound strategies have previously been adopted in the Li-Stephens model within the Eagle2 [16, 17] and fastLS [18] algorithms. The algorithm used in Threads incorporates some of the algorithmic innovations presented in fastLS. The output of the branch-and-bound steps is a set $\Omega$ of *path segments* $\omega$, each consisting of

- a start site $m_\omega \in \{1, \ldots, M\}$,
- a threading target $n_\omega \in \{1, \ldots, N\}$,
- if $m_\omega > 0$, a *traceback* segment $\omega' \in \Omega$ satisfying $m_{\omega'} < m_\omega$.

A full path through $\mathcal{H}$ can thus be constructed by starting at any path segment and following the traceback segments until a segment with $m_\omega = 0$ is found. We denote a path ending with $\omega$ as $\mathcal{P}(\omega)$ and by $s(\omega)$ the negative log-likelihood (penalty) of that path. The penalty for the partial path up to site $m$ is denoted by $s(\omega, m)$.

The set $\Omega$ contains exactly $N$ path segments $\omega_1, \ldots, \omega_N$ with $n_{\omega_n} = n$ classified as *active*, with the rest denoted *inactive*. If the active segments all satisfy the property that $\mathcal{P}(\omega_n)$ is a Li-Stephens-optimal path through $\mathcal{H}$ conditional on the path ending at $n$, the segment set is said to be *complete*. In this case, the path $\omega_n$ with the minimum penalty from among the active segments is a Viterbi path through $\mathcal{H}$.

Given a genotype vector $h_{m+1}$ for site $m+1$ and a complete segment set $\Omega^{(m)}$ for the partial panel $\mathcal{H}^{(m)}$ consisting of sites 1 through $m$, the branching step constructs a segment set $\Omega^{(m+1)}$ containing $\Omega^{(m)}$ that is complete for the partial panel $\mathcal{H}^{(m+1)}$. By contrast, the bounding step takes a complete segment set $\Omega^{(m)}$ and gives a smaller set $\Omega_*^{(m)} \subseteq \Omega^{(m)}$ that is also complete. Following the terminology of [18], we can say each segment in $\Omega^{(m)}$ is *undercut* by a segment in $\Omega_*^{(m)}$. The set $\Omega^{(0)}$, which contains only $N$ active segments $\omega_1, \ldots, \omega_N$ each starting at site 0 with a penalty of 0, is trivially complete for $\mathcal{H}^{(0)}$. By inductively applying the branching step at each site, we eventually arrive at a full Viterbi path. These steps are described in the following two theorems and in Figure 1.

**Theorem 1** (Branch). *Suppose $\Omega^{(m)}$ is a complete segment set for $\mathcal{H}^{(m)}$, and let $\omega' \in \Omega^{(m)}$ be a path segment such that $\mathcal{P}(\omega')$ is a Viterbi path through $\mathcal{H}^{(m)}$. Denote by $\rho$ and $\rho_c$ the penalties for recombination and no recombination between sites $m$ and $m+1$, respectively. Define*

$$\Omega' := \{\omega(m+1, n, \omega') \mid s(\omega_n, m) + \rho_c > s(\omega', m) + \rho\},$$

*where $\omega_n$ are the active segments in $\Omega^{(m)}$. Then, the segment set*

$$\Omega^{(m+1)} := \Omega^{(m)} \cup \Omega'$$

*obtained by replacing the active segments of $\Omega^{(m)}$ with those from $\Omega'$ where possible, is a complete segment set for $\mathcal{H}^{(m)}$.*

*Proof.* Fix $n \in \{1, \ldots, N\}$ and let $s_{m+1}$ denote the penalty of a Li-Stephens-optimal path through $\mathcal{H}^{(m+1)}$ ending in $n$. Let $\mu$ be the match/mismatch penalty for the target haplotype $g$ and $h_n$ at $m + 1$. Then, by definition,

$$s_{m+1} \leq \min(\{s(\omega_n) + \rho_c + \mu,\ s(\omega') + \rho + \mu\}).$$

If $s_{m+1} < s(\omega_n)+\rho_c+\mu$, then any path with penalty $s_{m+1}$ ending at haplotype $n$ must recombine between $m$ and $m+1$. By Proposition 1, $s_{m+1} = s(\omega') + \rho + \mu$. Similarly, if $s_{m+1} < s(\omega') + \rho + \mu$, then any path with score $s_{m+1}$ ending at haplotype $n$ does not recombine between sites $m$ and $m + 1$, giving $s_{m+1} = s(\omega_n) + \rho_c + \mu$. Thus, it follows that $s_{m+1} = \min(\{s(\omega_n)+\rho_c+\mu,\ s(\omega')+\rho+\mu\})$, which is exactly the penalty for the active state for haplotype $n$ in $\Omega^{(m+1)}$. □

Starting from $\Omega^{(0)}$, sequential application of this branching step generates $N$ distinct tree structures, with segments for nodes and traceback pointers as edges. At each step, the complete segment set grows by at most $N$ elements, resulting in a time and memory complexity of $O(NM)$. However, new information is stored only in the case of inferred recombination events and, as previously observed, recombination events in Viterbi paths through realistic genetic data are very rare. Thus, in practice, the memory usage grows substantially slower than $O(MN)$. In cases where there is rapid growth in memory usage or other constraints, even this level of memory usage may not be computationally feasible. For this reason, we include a step to prune the complete segment set and limit the number of segments that need to be stored in memory at any one time.

**Theorem 2** (Bound). *Let $\Omega$ be a complete segment set through $\mathcal{H}$ with active segments $\omega_1, \ldots, \omega_N$. Define $\Omega_* \subseteq \Omega$ as the segment set $\bigcup_{n=1}^{N} \mathcal{P}(\omega_n)$, which contains those segments*

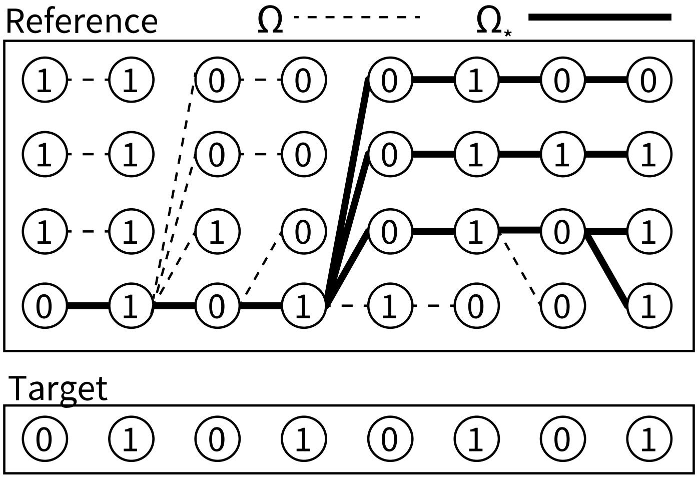

Figure 1: Diagram of the Threads-Viterbi algorithm. The full set of expanded segments $\Omega$ (both dashed and thick lines) is replaced with the subset $\Omega_*$ (thick lines). In this instance, $|\Omega| = 13$ and $|\Omega_*| = 5$. A unique path of lowest penalty given the target sequence can be traced back for each of the four reference sequences.

*that are part of a traceback path from one of the active segments in $\Omega$. Then, $\Omega_*$ is also a complete segment set through $\mathcal{H}$.*

*Proof.* By Theorem 1, it is sufficient to expand only the active segments during the branching step and replace them with new segments. This means that the completeness condition depends solely on the active segments. By the definition of the segment sets, and to allow for Viterbi paths to be traced back, it is necessary to include those segments that are part of the traceback paths. The set $\Omega_*$ contains exactly those segments. □

We apply the bounding step at regular intervals throughout the inference process. We use either fixed intervals or follow a heuristic that limits the growth rate of the total number of segments. In the latter case, we fix a starting value (in Threads, we use $B_0 := 10 \cdot N$), and then, if $|\Omega^{(m)}| > B_0$, we prune $\Omega^{(m)}$ to $\Omega_*^{(m)}$. If again $|\Omega^{(m')}| > B_0$ for some $m' > m$, we continue with a new threshold of $B_1 := 2B_0$ if $m' - m < 30$, otherwise we continue with the threshold of $B_0$. This procedure heuristically balances the number of pruning operations against the risk of memory spikes due to rapid accumulation of segments. We never need to store either

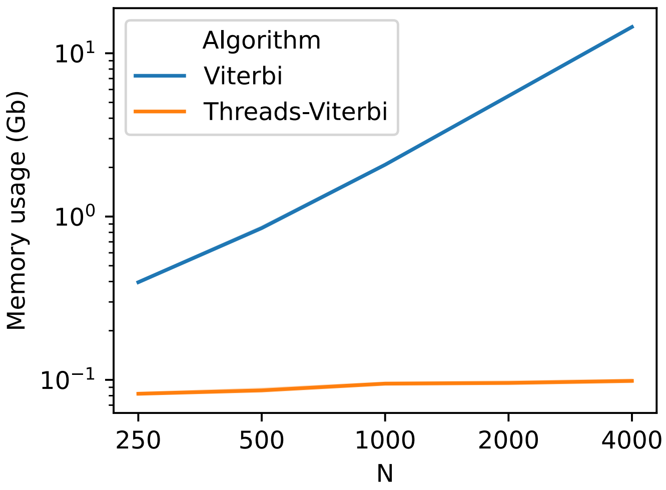

Figure 2: Memory usage of the Threads-Viterbi algorithm and the classical Viterbi algorithm on simulated genotype data. Sequences of 5MB length were simulated under a European demographic model. In this analysis we applied the bounding step of Threads-Viterbi every 10 sites.

the full genotype matrix in memory or any $N \times M$ probability matrix.

In sum, we obtain an algorithm with $O(NM)$ runtime and, on average, $O(N)$ memory usage. Pseudocode is provided in Algorithm 2 and a diagram of the algorithm is given in Figure 1. In practice, the penalty function $s(\cdot)$ is never evaluated on a full or partial path, instead penalties are updated as attributes of the active segments. In Algorithm 2, we use the penalty function $s$ only to simplify notation. A comparison between memory usage of the Threads-Viterbi algorithm and the classical Viterbi is shown in Figure 2.

The algorithm we have described thus far finds the Viterbi path through a single $N \times M$ genotype matrix. To compute a full set of threading instructions, however, we must infer such a path for each sample $n$, using a reference panel comprising samples $1, \ldots, n-1$. Using the haplotype matching step described above, we obtain a panel of size approximately $L \times M$ for each sample $n$, for some $L \ll N$. Crucially, the output of each hidden Markov model does not depend on any of the other hidden Markov models, meaning that we can safely run the $N$ instances of Threads-

**Algorithm 2** Threads-Viterbi

**Input:** Genotype matrix $X \in \{0,1\}^{M \times N}$, target genotype $g$, genetic distances $c$, physical distances $d$, and expected coalescence height $t$.

**Output:** A Viterbi path through $X$ for $g$.

$\Omega \Leftarrow \{\omega_0, \dots, \omega_{N-1}\}$ ▷ Initialize the segment set with empty segments
$\omega^* \Leftarrow \omega_0$ ▷ Initialize the best segment
**for** $m = 0, \dots, M-1$ **do**
  $\mu_c = 2 * \mu * d_m * t$ ▷ Mismatch penalties for site $m$
  $\mu = -\log(1 - e^{-\mu_c})$
  $\rho_c = 0.02 * c_m * t$ ▷ Recombination penalties for site $m$
  $\rho = -\log(1 - e^{-\rho_c})$
  $\omega' \Leftarrow \omega^*$
  $s' \Leftarrow s(\omega^*, m-1) + \mu + \rho$
  **for** $i = 0, \dots, N-1$ **do** ▷ Perform the branching step
    $\mu_i \Leftarrow \mu$ if $X_{m,i} \neq g_m$, else $\mu_c$
    $s' \Leftarrow \min(s(\omega_i, m-1) + \rho_c, s(\omega^*, m-1) + \rho) + \mu_i$
    **if** $s(\omega_i, m-1) + \rho_c > s(\omega^*, m-1) + \rho$ **then**
      Add new segment $\omega(m, i, \omega^*)$ to $\Omega$.
      Set the new segment to be the active segment $\omega_i$ for $i$.
    **end if**
    **if** $s' < s(\omega_i, m)$ **then**
      $\omega' \Leftarrow \omega_i$
      $s' = s(\omega_i, m)$
    **end if**
  **end for**
  $\omega^* \Leftarrow \omega'$
  **if** $|\Omega| > B$ **then** ▷ Perform the bounding step if $|\Omega|$ exceeds a threshold
    $\Omega' \Leftarrow \{\omega_0, \dots, \omega_{N-1}\}$
    **for** $i = 0, \dots, N-1$ **do**
      $\omega \Leftarrow \omega_i$
      **while** $\omega$ has a starting site $> 0$ **do**
        $\omega \Leftarrow \text{traceback}(\omega)$
        Add $\omega$ to $\Omega'$
      **end while**
    **end for**
    $\Omega \Leftarrow \Omega'$
  **end if**
**end for**
$\mathcal{P} \Leftarrow \{\omega^*\}$ ▷ Final traceback step
$\omega \Leftarrow \omega^*$
**while** $\omega$ has a starting site $> 0$ **do**
  $\omega \Leftarrow \text{traceback}(\omega)$
  Add $\omega$ to $\mathcal{P}$
**end while**
**return** $\mathcal{P}$

Viterbi in parallel. In practice, we divide the $N$ HMMs evenly among available CPU cores. This means we can complete the Viterbi inference step for all samples using only a single pass through the data per core.

Given $L$ haplotype matches per sample, threading targets can thus be obtained in $O(MLN/N_{\text{CPU}})$ time and $O(LN)$ memory on average, streaming the genotypes from disk only once for the haplotype matching and once for the Threads-Viterbi algorithm. In comparison, $O(MN^2)$ time and memory are required to run the same inference without haplotype matching and using a classical HMM for the Li-Stephens inference. Even so, the Threads-Viterbi step is still the most computationally intensive part of the Threads algorithm.

The Viterbi paths may be interpreted as a map $[L_1, L_2] \to \{1, \ldots, n-1\}$ for each $n \leq N$, representing a threading target for each sample at each site. To complete the ARG inference, a third and final pass through the genotypes is required to assign coalescence times to each of these inferred threading targets. The next chapter describes and motivates the segment-based coalescent modeling used in the last step of Threads.

### 4 Dating path segments

In this section, we describe the modeling used to assign coalescence times to each of the segments inferred by the Threads-Viterbi algorithm. These dated Viterbi segments constitute the threading instructions needed to assemble the ARG. To efficiently estimate the ages of these segments, we model them as identical-by-descent (IBD) regions that are shared by the target sample and the set of closest cousins. We further assume these segments to be independent. Together with the Threads-Viterbi step, this dating method forms a change-point detection algorithm, partitioning a stretch of genome into segments of constant height and threading target. This type of model is sometimes referred to as a product partition model [2].

We define a genomic region delimited by $[L_1, L_2]$ to be *identical by descent* (IBD) for a pair of sequences if their most recent common ancestor is constant along the region. We say $[L_1, L_2]$ is an *IBD segment* if it cannot be extended in either direction while remaining IBD. IBD segments have a fixed *height*, or *age*, measured by the age of the common ancestor, in generations. Given the length and the number of heterozygous sites along the IBD segment, we use these modeling assumptions to estimate the segment age. In addition, we use an estimated demographic history for the pair of analyzed sequences, measured by the effective population size $N_e(t)$, as a prior estimate of the segment age. In the following, we derive the formulas used to estimate coalescence times.

We consider an IBD segment of length $l_{\text{cM}}$ in centimorgans (cM) and $l_{\text{bp}}$ in base-pairs (bp), and begin by assuming a fixed mutation rate $c$ and a constant coalescence rate $\gamma$. We model the number of heterozygous sites along the segment to follow a Poisson distribution with rate $\mu = 2 \cdot c \cdot l_{\text{bp}}$. Given the starting coordinate of the segment, we model the recombination process according to the sequentially Markovian coalescent (SMC) model [20], such that the segment's length follows an exponential distribution with rate $\rho = 2 \cdot 0.01 \cdot l_{\text{cM}}$. Under these assumptions, we derive several estimates of increasing complexity and accuracy for the segment age $t$.

#### 4.1 Maximum likelihood estimators

We first disregard the coalescence rate $\gamma$ and focus on likelihood functions for $t$ using $\rho$ and $\mu$. Ignoring mutations, under the SMC model, the length (in Morgans) of an IBD segment follows an exponential distribution with a rate of $1/2t$. We derive a likelihood $\ell(\rho|t) = 2te^{-t\rho}$, which has derivative $2e^{-t\rho}(1 - t\rho)$ with respect to $t$ and a maximum at $t = 1/\rho$.

To incorporate mutations into the likelihood function, we denote the number of heterozygous sites on the segment as $m$ and express

$$\ell(\rho, m|t, c) = \ell(\rho|t, c)\ell(m|\rho, t, c) = 2te^{-t\rho}\frac{(t\mu)^m}{m!}e^{-t\mu}.$$

Differentiating with respect to $t$, we find that this expression has a maximum at $t = \frac{m+1}{\rho+\mu}$. In what follows, we will drop the mutation rate $c$ from the notation.

#### 4.2 Bayesian estimators

We now place a prior $\pi$ on $t$ and compute $\mathbb{E}[t|\rho, m]$, assuming $\pi(t) \sim \text{Exp}(\gamma)$. This approach is distinct from using the prior $\underline{\pi}(t) \propto t\pi(t)$ which models coalescence at each site rather than each segment, and accounts for the inverse correlation between segment height and length [6, 8, 12, 21].

As above, we start with a point estimate for $t$ that ignores mutations. We observe that

$$p(t|\rho) \propto \pi(t)p(\rho|t) = 2\gamma t e^{-t(\rho+\gamma)},$$

an Erlang-2 distribution with rate $\rho + \gamma$ and mean $\frac{2}{\rho+\gamma}$ [8, 22]. Adding mutations, we can similarly write

$$p(t|\rho, m) \propto \pi(t)p(m|\rho, t)p(\rho|t) = 2\frac{\gamma\mu^m}{m!}t^{m+1}e^{-t(\rho+\mu+\gamma)}$$

to obtain an Erlang-$(m + 1)$ distribution with rate $\rho + \mu + \gamma$ and mean $\frac{m+2}{\rho+\mu+\gamma}$. We refer to the estimators $\mathbb{E}[p(t|\rho, m)]$ and $\mathbb{E}[p(t|\rho)]$ as the *Bayesian estimators*.

#### 4.3 Piecewise-constant demographic models

We now extend the estimators above to account for variable demographic models. In problems dealing with variable demographic models, a standard simplification is to assume that the effective population size $N_e(t)$ is piecewise constant [14]. Under this assumption, we write $0 = T_0 < T_1 < \cdots < T_K = \infty$ and assume $N_e(t) \equiv N_e^{(k)}$ for all $t \in [T_k, T_{k+1})$. Similarly, we obtain piecewise-constant coalescence rates $\gamma_k = 1/N_e^{(k)}$ within each interval. Writing $\Delta_k = T_{k+1} - T_k$ for $k = 0, \ldots, K-1$, the prior becomes $\pi(t) = \gamma_k e^{-\sum_{j=0}^{k-1}\Delta_j\gamma_j - (t-T_k)\gamma_k}$ for $t \in [T_k, T_{k+1})$. In this case, there is no compact description of the distributions $p(t|\rho)$ and $p(t|\rho, m)$, so instead we compute the expectations directly.

As before, we initially ignore the number of heterozygous sites. This gives us the ratio of integrals

$$\mathbb{E}[t|\rho] = \int_0^\infty tp(t|\rho)\mathrm{d}t = \frac{\int_0^\infty t\pi(t)p(\rho|t)\mathrm{d}t}{\int_0^\infty \pi(t)p(\rho|t)\mathrm{d}t}.$$

The numerator has solution

$$\int_0^\infty t\pi(t)p(\rho|t)\mathrm{d}t = \sum_{k=0}^{K-1}\int_{T_k}^{T_{k+1}} t\gamma_k e^{-\sum_{j=0}^{k-1}\Delta_j\gamma_j-(t-T_k)\gamma_k}2te^{-t\rho}\mathrm{d}t$$
$$= \sum_{k=0}^{K-1}\gamma_k e^{-\sum_{j=0}^{k-1}\Delta_j\gamma_j+T_k\gamma_k}2\int_{T_k}^{T_{k+1}} t^2 e^{-t(\rho+\gamma_k)}\mathrm{d}t$$
$$= \sum_{k=0}^{K-1}\gamma_k e^{-\sum_{j=0}^{k-1}\Delta_j\gamma_j+T_k\gamma_k}\frac{4}{(\rho+\gamma_k)^3}$$
$$\times\Big[P(3,(\rho+\gamma_k)T_{k+1}) - P(3,(\rho+\gamma_k)T_{k+1})\Big],$$

where

$$P(a,z) = \frac{\gamma(a,z)}{\Gamma(a)} = \frac{1}{\Gamma(a)}\int_0^z t^{a-1}e^{-t}\mathrm{d}t$$

is the normalized lower incomplete gamma function. The denominator has a similar expression, and together they provide the point estimate

$$\mathbb{E}[t|\rho] = \frac{\sum_{k=0}^{K-1}\gamma_k e^{-\sum_{j=0}^{k-1}\Delta_j\gamma_j+T_k\gamma_k}\frac{2}{(\rho+\gamma_k)^3}\Big[P\big(3,(\rho+\gamma_k)T_{k+1}\big) - P\big(3,(\rho+\gamma_k)T_{k+1}\big)\Big]}{\sum_{k=0}^{K-1}\gamma_k e^{-\sum_{j=0}^{k-1}\Delta_j\gamma_j+T_k\gamma_k}\frac{1}{(\rho+\gamma_k)^2}\Big[P\big(2,(\rho+\gamma_k)T_{k+1}\big) - P\big(2,(\rho+\gamma_k)T_{k+1}\big)\Big]}. \tag{1}$$

Using the same approach, it is straightforward to add mutations to the model to get the estimate

$$\mathbb{E}[t|\rho,m] = \frac{\sum_{k=0}^{K-1}\gamma_k e^{-\sum_{j=0}^{k-1}\Delta_j\gamma_j+T_k\gamma_k}\frac{\mu^m(m+2)}{\lambda_k^{m+3}}\Big[P\big(m+3,\lambda_k T_{k+1}\big) - P\big(m+3,\lambda_k T_k\big)\Big]}{\sum_{k=0}^{K-1}\gamma_k e^{-\sum_{j=0}^{k-1}\Delta_j\gamma_j+T_k\gamma_k}\frac{\mu^m}{\lambda_k^{m+2}}\Big[P\big(m+2,\lambda_k T_{k+1}\big) - P\big(m+2,\lambda_k T_k\big)\Big]}, \tag{2}$$

where $\lambda_k = \rho + \mu + \gamma_k$. This is the estimate used by Threads to infer coalescence times in sequencing data. In inference from sparse sequences such as genotyping array data, we rely on the mutation-free formula in Equation 1.

### 5 Modeling limitations

In this section we discuss several caveats to the IBD-based model used in Threads to estimate the age of Viterbi segments. First, Viterbi segments do not necessarily coincide with IBD segments. In Figure 3, we show in simple toy examples how Viterbi paths, even when inferred perfectly, may both over- and underestimate the length of the IBD segments they reflect. We can quantify the frequency of these events in a simple simulation as follows. Fix a leaf $l$ in a simulated ARG, with most recent ancestor $a_1$ in a neighborhood left of some fixed site $x$ and a different most recent ancestor $a_2$ to the right of $x$. The closest cousins of $l$ at either side of $x$ belong to different IBD segments, and we want to quantify to what extent these coincide with Viterbi segments. If the leaf sets subtended by $a_1$ and $a_2$ (excluding $l$) have a non-empty intersection, then this recombination event is not detectable by the Viterbi algorithm and we overestimate the length of the underlying IBD segment. This case is illustrated in Figure 3c. If the leaf sets are disjoint, but $a_2$ is an ancestor of $a_1$ in a neighborhood to the left of $x$ or $a_1$ an ancestor of $a_2$ to the right of $x$, the IBD segments overlap and we underestimate their length by choosing a single breakpoint. We call these events "left" and "right" overlaps and give an illustration in Figures 3a and 3b. If none of these conditions are satisfied we say the Viterbi segment coincides with the IBD segments.

To quantify the frequency of each of these events we traversed simulated ARGs of varying size, annotating changes in the most recent common ancestor above a fixed leaf, as presented in Figure 4. We observed that the rate at which Viterbi segments coincided with IBD remained stable at $\sim 40\%$, while the rate at which this approach overestimated segment lengths tended to $0$ as $N$ increased. This occurs because overestimates can only happen when a recombination event results in the same genealogical closest cousin belonging to two adjacent IBD segments. As the ARG grows, the proportion of volume taken up by the clade subtending the fixed leaf $l$ and its closest cousins is diminished, making this type of event less likely. Overall, these results suggest that the heuristic approach of dating Viterbi segments by treating them as pairwise IBD regions might underestimate the segment length and thus overestimate their height.

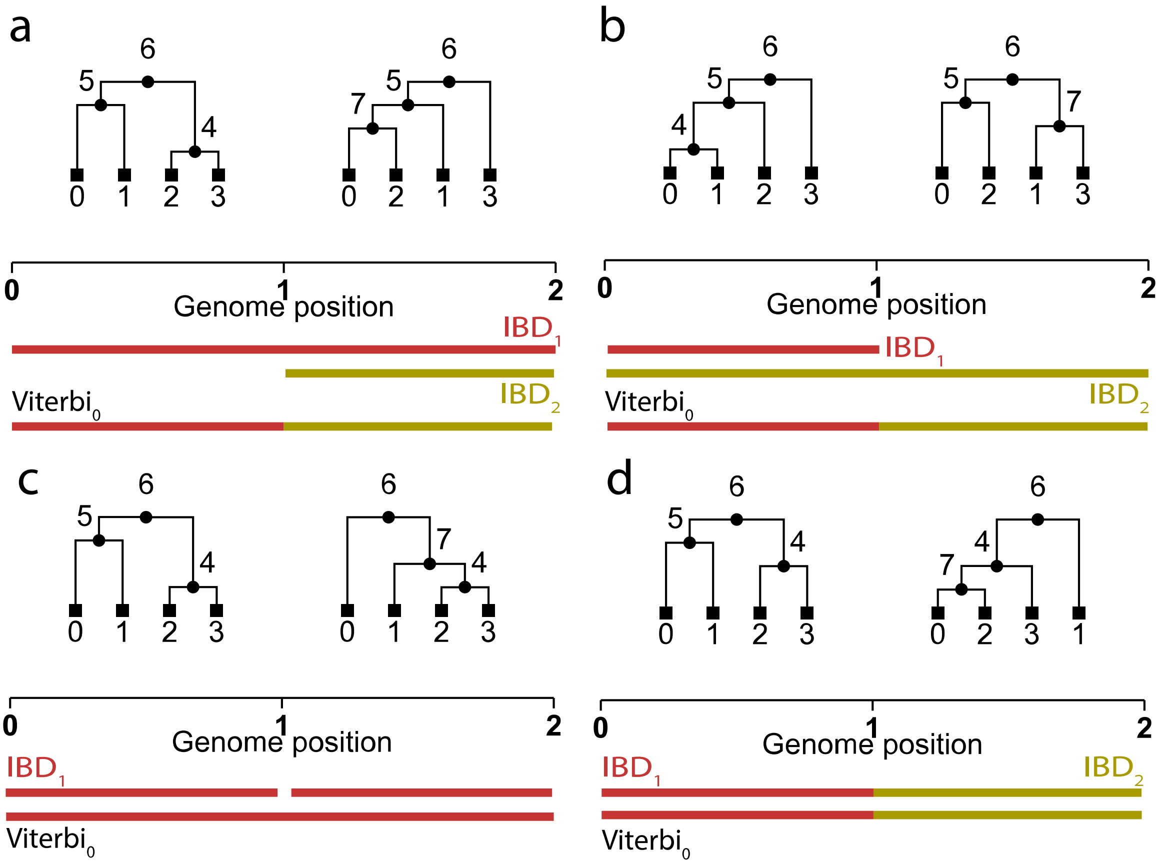

Figure 3: Illustration of how Viterbi segments might differ from IBD segments. Colored segments show Viterbi and IBD segments for the leaf labeled 0. In each panel, a unique Viterbi path can be found for leaf 0 through leaves 1 and 2. In panels **a** and **b**, the IBD segment subtended by the ancestor labeled 5 overlaps the next or the previous IBD segment. We call these events "right" and "left" overlaps, respectively. In panel **c**, a single, observed Viterbi segment overlaps two distinct IBD segments, overestimating the actual segment length. Finally, panel **d** shows a scenario where the IBD and Viterbi segments coincide.

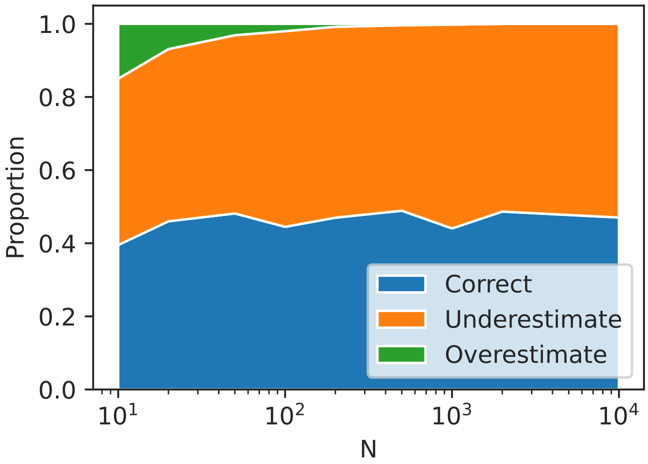

Figure 4: Concordance of IBD segments and Viterbi segments in simulated ARGs under a European demographic model. Here, we fix a leaf and trace it along the genome until the first change in the most recent common ancestor. We repeat this experiment, varying the random seed and the total number of leaves $(N)$ from $10$ to $10{,}000$. We then estimate the probability of observing each of the recombination patterns from Figure 3, which we label as "underestimate" (Fig 3a-b), "overestimate" (Fig 3c) and "correct" (Fig 3d).

In practice, noisy inference of Viterbi segments will also affect their length. In real data sets, the algorithm tends to infer segments that are longer and fewer in number than the true underlying segments, especially for smaller sample sizes, as shown using simulations in Figure 5. For small sample sizes, Threads output less than half the true number of segments for ARGs on tens of samples, with the difference diminishing as sample sizes approach the tens of thousands. Inferring fewer and longer segments also leads to underestimating their height.

As a result, the IBD-based approach used within Threads relies on modeling simplifications that lead to both over- and underestimation of segment ages. In practice, we have observed this approach to be computationally fast and lead to reasonable accuracy. We discuss some ways to improve segment dating in the small sample regime below, leaving more sophisticated methods to model Viterbi segment lengths as future work.

The second main caveat to pairwise IBD-based estimates of coalescence times is related to the choice of coalescence time prior. In the segment age estimates derived in Equation 2, it is assumed that the two sequences coalesce with a pairwise population-based rate of $\gamma$. However, this approach ignores the fact that at each site the IBD segments we aim to date represent the first coalescence event of the sequence being threaded with any of the $n-1$ sequences that are already present in the ARG. A better model of the first coalescence time of sample $n$ comes from using a variable coalescence rate of $\gamma(t) \cdot L_{n-1}(t)$, where $L_{n-1}(t)$ counts the number of lineages in the ARG on $n-1$ leaves at time $t$. The distribution of the random variable $L_{n-1}(t)$ has a closed-form expression, obtained by marginalizing over its possible values, but takes $O(n)$ time to evaluate.

Using the exact prior, evaluating an estimator like the one in Equation 2 would take $O(CN)$ time, where $C$ is the number of intervals in which the demographic model is discretized. This would lead to a time complexity of $O(CN^2)$ for the whole ARG-inference process, which is prohibitively expensive. Again, we found the heuristic use of a pairwise coalescence rate $\gamma(t)$ to be sufficiently accurate, and opted to leave the use of a more accurate and computationally scalable prior as future work. To verify this, we evaluated the accuracy of estimation of the time to the most recent common ancestor (TMRCA) between the sample being threaded

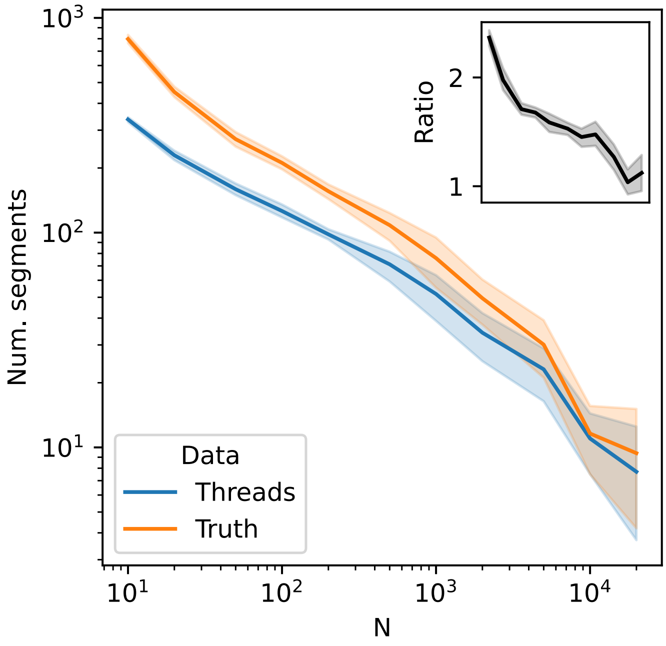

Figure 5: The number of segments inferred by Threads compared with the true number of underlying IBD segments. Here, we simulate ARGs for $N$ ranging from $10$ to $20{,}000$ under a European demographic model with fixed mutation rate $\mu = 1.65 \times 10^{-8}$ across a $20$ megabase region for $10$ random seeds. We infer an ARG from the observed genotypes using Threads and count the number of distinct segments for the last sample to be threaded. We then compare this with the true number of segments from the simulated ARGs for that sample.

and its inferred closest cousin, and of the actual first coalescence time. Results of these simulations as shown in Figure 6 for ARGs comprising $100$, $1{,}000$, and $10{,}000$ samples. Correlation estimates (Pearson's $r$) are sensitive to outliers and fluctuated significantly, but we observed robust rank correlation (Spearman's $r$). Since threading-based ARG inference relies on the relative timing of segments to estimate topology, a high rank correlation of estimated coalescence times suggests good topological accuracy.

The final simplifying assumption that we highlight is the independence of segment heights. In the SMC and related frameworks it is standard to model the coalescence times of adjacent sites using a conditional distribution $p(t_1|t_2)$, where the adjacent coalescence times $t_1$ and $t_2$ are not independent, and several approaches to computing $p(t_1|t_2)$ exist [10, 19, 20]. Threads instead implements a change point detection algorithm, where the change points are inferred jointly in the Viterbi step, but their age inferred independently in a second step (also see [6, 12]).

Each of the simplifications discussed above brings substantial computational benefits. The inference of fewer segments leads to ARGs containing fewer edges and nodes compared to the threading approach used in ARG-Needle [26], significantly reducing the memory consumption and computational costs of downstream analyses. The simplified demographic prior allows estimating the height of each segment in $O(C)$ time for $C$ discretized time intervals in the demographic model. The independence of segment heights also allows parallelizing this computation, as done for the Li-Stephens inference step. Overall, we found Threads to perform well across several benchmarks and to allow scaling of ARG inference to very large sample sizes, suggesting that the computational benefits gained from applying these modeling simplifications do not result in a substantial loss of accuracy.

### 6 Inference heuristics

#### 6.1 Li-Stephens parameters

Mismatch and recombination rates for the Li-Stephens model are usually either estimated numerically or modelled using coalescent arguments that rely on panel

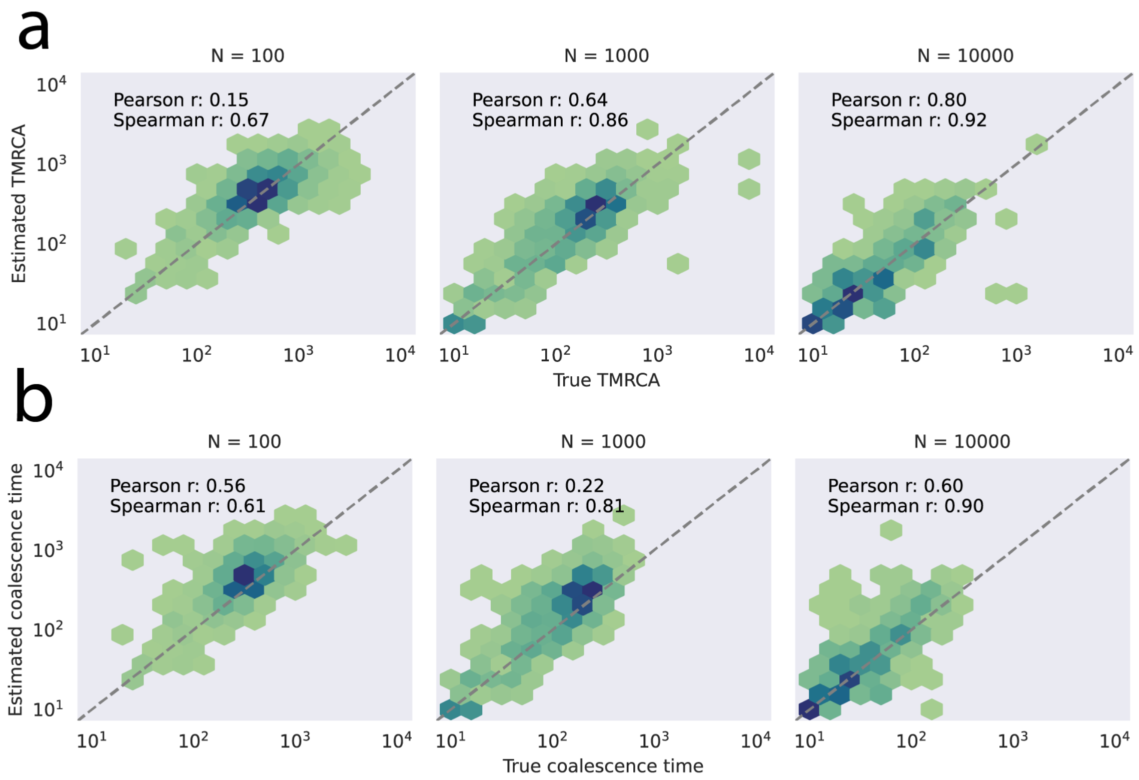

Figure 6: Accuracy of TMRCA inference using segment-based dating. We simulated ARGs for $N$ ranging from $100$ to $10{,}000$ under a European demographic model with fixed mutation rate $\mu = 1.65 \times 10^{-8}$ across a $20$ megabase region for $10$ random seeds. We then inferred ARGs for the observed genotypes using Threads. In **a**, we compare the coalescence times of the last $5$ samples to be threaded into the ARG to the actual TMRCA of that sample and the closest cousin as inferred by Threads. Coalescence times are observed at $20$ evenly spaced points between coordinates $5{,}000{,}000$ and $15{,}000{,}000$, allowing sufficient burn-in for segment dating and closest cousin inference. In **b**, we compare the inferred coalescence times of the last $5$ samples to be threaded into the ARG with their actual coalescence times, at the same base-pair coordinates as in panel **a**.

size, empirical recombination and mutation rates, and demographic history. For Threads, the panel size changes at each iteration, making numerical estimates unfeasible, so we instead rely on modeling.

Given a tree on $N$ leaves under the standard coalescent [13], the expected time to the first coalescence event for a leaf is $2/N$. The coalescence time of the $N$-th sequence being threaded and the $N-1$ sequences previously threaded are jointly distributed as a coalescent tree on $N$ leaves. Thus, the expected coalescence time of the $N$-th sequence into an ARG also becomes $2/N$. In a constant-sized population of $N_e$ haploids, this expected coalescence time becomes $2N_e/N$. For two sequences coalescing $2N_e/N$ generations ago, the distance in centimorgans to the next recombination event follows an exponential distribution with rate $0.01 \cdot 2N_e/N$. This quantity is used set the recombination probabilities of $1 - e^{0.01 \cdot l \cdot 2N_e/N}$ in the Li-Stephens model where $l$ measures the distance between sites in centimorgans.

We extend this model to piecewise-constant demographic models, which can be done by applying a piecewise linear monotonic re-scaling of time. A demographic model with variable population sizes, $N_e(t)$, gives a mapping from absolute time, measured in generations, to standard coalescence time via $t \mapsto \int_0^t N_e(s)^{-1} \mathrm{d}s$.

For a piecewise-constant demographic model with time breakpoints $0 = t_1 < t_2 < \cdots < t_K$ and effective haploid population sizes $N_1, N_2, \ldots, N_K$, we compute the expected coalescence times as follows. We first map each of the breakpoints to coalescent time using the map $t_i \mapsto \tau_i := \tau_{i-1} + (t_i - t_{i-1})/N_{i-1}$ for $i > 1$, with $\tau_1 = 0$. To get the expected coalescence time for $N$ leaves under this demographic model, we then linearly interpolate between the values $\tau_i$ and then change coordinates back to time in generations using the inverse map. In practice, this means finding the highest $i$ such that $\tau_i \leq 2/N$, giving the expected coalescence time $T_N := t_i + (2/N - \tau_i)N_i$. This leads to a recombination probability of $1 - e^{-0.01 l \cdot 2T_N/N}$ for sites at a distance of $l$ centimorgans under a tree on $N$ leaves and a mismatch probability of $1 - e^{-\mu l' 2T_N/N}$ on a tract of genome of length $l'$ base-pairs under a constant per-site-per-generation mutation rate of $\mu$.

#### 6.2 ARG inference from genotyping array data

In several cases, sequencing data is not available, and analyses instead rely genotyping array data, often imputed using a reference panel of sequenced individuals. Imputation may not always be available, or, in cases where the ancestry of the analyzed samples is poorly represented in the reference panel, it may yield noisy genotype data, particularly for rare variants. The ARG-Needle algorithm [26] was designed for the inference of ARGs from genotyping array data, and has been leveraged to complement genotype imputation in the analysis of rare variation in complex traits. Here, we describe how Threads can be adjusted for the analysis of SNP array data sets.

Missing variants in genotyping array data sets cause the misspecification of the emission probabilities for the Li-Stephens HMM. The variants present on these arrays are not randomly ascertained and are biased towards more common variation. The ascertained sequentially Markovian coalescent (ASMC) algorithm [23] uses approximate frequency-based modeling to account for these biases, which is built on the conditional site frequency spectrum (CSFS) [25]. Threads relies on a more heuristic approach, based on artificially inflating the rate of recombination events.

If the probability of recombination between two sites in the Li-Stephens model is $p$, then the transition penalty in the HMM is $-\log(p/N)$. When modeling sparse sites from genotyping array data in Threads, we instead use a transition penalty of $-\log(p)$. This heuristic is applied in cases where common variants are expected to be overrepresented in the subset of variants that have been ascertained. The lower transition penalty encourages recombination in this low-signal regime. Mutation penalties in the sparse model are kept identical to the full whole-genome model.

This model differs from other array-based Li-Stephens implementations, such as Beagle 5.4 [4] and IMPUTE5 [24]that use a fixed mismatch probability of $0.0001$ and recombination rates identical to the full model. However, these rates are optimized for biobank-scale panel sizes and not guaranteed to generalize to the whole process of ARG inference. In our experiments we find that Threads is relatively robust to the choice of Li-Stephens parameters, with slightly more accurate results observed using the heuristic sparse-model parameters described above. However, we caution that the exact choice of parameters is dataset-specific and may not fit well in other data sets.

#### 6.3 Small-$N$ inference

As previosuly discussed, Viterbi paths through a reference panel do not necessarily correspond to underlying IBD segments and the difference is especially pronounced in smaller sample sizes. Accurate threading of small sample sizes is crucial to get high-quality inference of deep-time coalescence events, so the samples that are initially threaded into the ARG have a disproportionate weight on the quality of the ARG at deep time scales.

To improve threading accuracy in small sample sizes, we implemented a Viterbi algorithm based on the pairwise sequentially Markovian coalescent (PSMC) [14] that we apply to any haplotype matching segment with more than 5 heterozygous sites inferred while threading the first 1,000 sequences. The algorithm identifies the most likely path through a set of discrete time states, and we interpret each jump in the PSMC path as a recombination event defining a new IBD segment. This operation breaks the input segment into one or more sub-segments that we each date independently using the same approach as in the base model.

In our experiments, we found that the addition of the PSMC step increased accuracy for the inference of deep coalescence times when inferring ARGs from dense marker sets, but led to limited gains in the analysis of genotyping array data. By default, this step is included in inference from dense marker ("sequencing mode") sets but omitted when Threads is run on genotyping array data ("array mode").

Using the modular nature of the threading instructions, it is also possible to *seed* Threads ARG inference with an independently inferred ARG, built using a small batch of sequences and applying methods which are less scalable but that rely on more accurate modeling of deep-time genealogical processes. This approach, however, requires further methodological development, as we observed that different calibration of coalescence times between different models can yield sub-optimal results.
