## Supplementary Figures for "A scalable approach for genome-wide inference of ancestral recombination graphs"

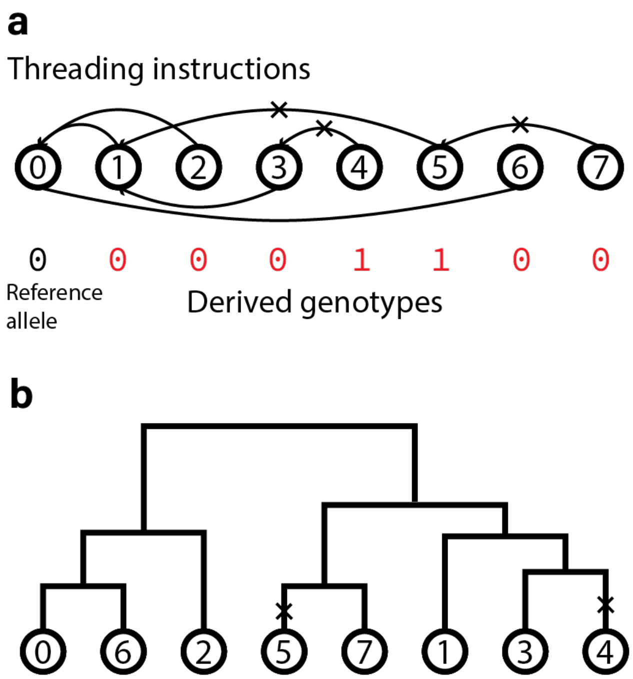


**Supplementary Figure 1. Relationship between threading instructions and genotype data. (a)** A cartoon showing the rooted, directed tree structure defined by threading instructions at a single site. Circled numbers represent indexed genomes, edges indicate threading partner relations and 'x' symbols indicate the two sequences are heterozygous at the site. Coalescence times, stored on each edge, are not represented in this image. Only the reference allele for the first sequence, with value 0, is stored in memory. To recover the genotype of other sequences, the tree is traversed towards the root and the reference allele flipped for each mutation ('x') encountered along the unique path to the root. For example, the unique path 7 → 5 →1 → 0 from sequence 7 to the root carries two mutations and thus the reference allele 0 is flipped two times, giving a genotype value of 0 for sequence 7. **(b)** A coalescence tree representing the same data in the example of panel (a)**.**

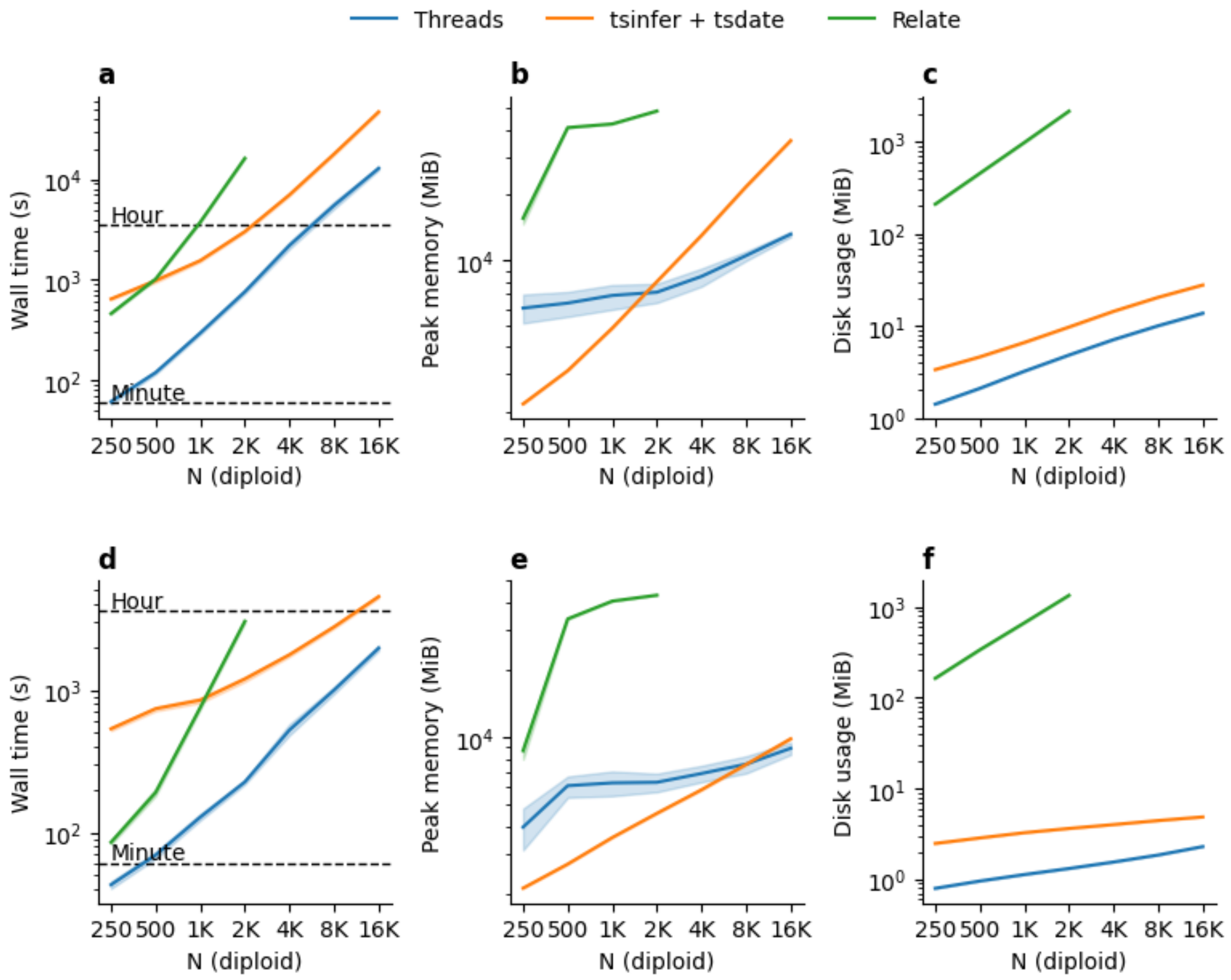


**Supplementary Figure 2. Computational performance on simulated sequencing data.** We measure runtime, peak memory usage, and disk space required for final output. **a-c**. Simulations under a European (CEU) demography. **d-f.** Simulations under a constant (Ne=10,000) demography. Shaded regions show bootstrap 95% confidence intervals across 15 random seeds.

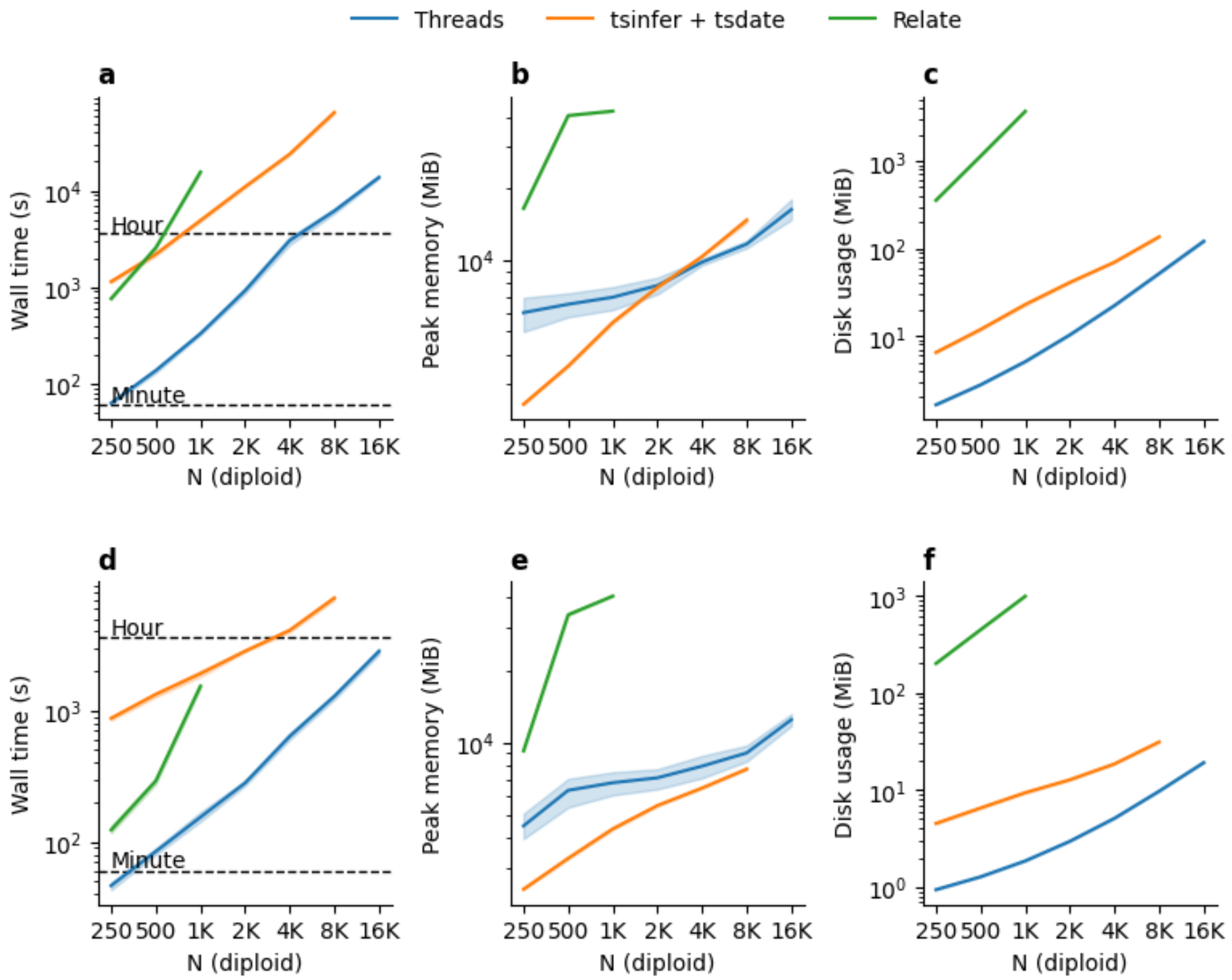


**Supplementary Figure 3. Computational performance on simulated sequencing data with a 0.1% error rate**. We measure runtime, peak memory usage, and disk space required for final output. **a-c**. Simulations under a European (CEU) demography. **d-f.** Simulations under a constant (Ne=10,000) demography. Shaded regions show 95% confidence intervals across 15 random seeds.

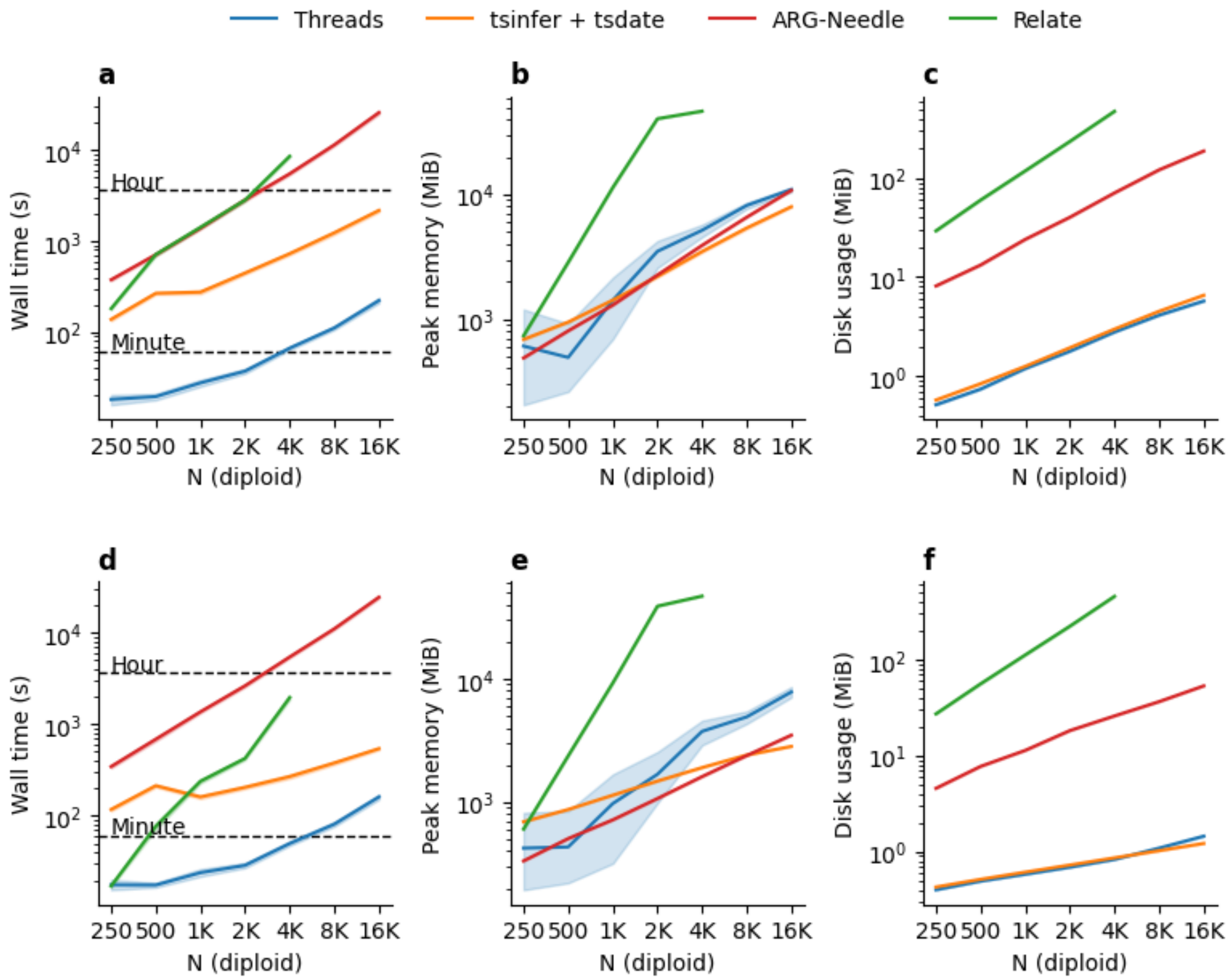


**Supplementary Figure 4. Computational performance on simulated genotyping array data**. We measure runtime, peak memory usage, and disk space required for final output. **a-c**. Simulations under a European (CEU) demography. **d-f.** Simulations under a constant (Ne=10,000) demography. Shaded regions show 95% confidence intervals across 15 random seeds.

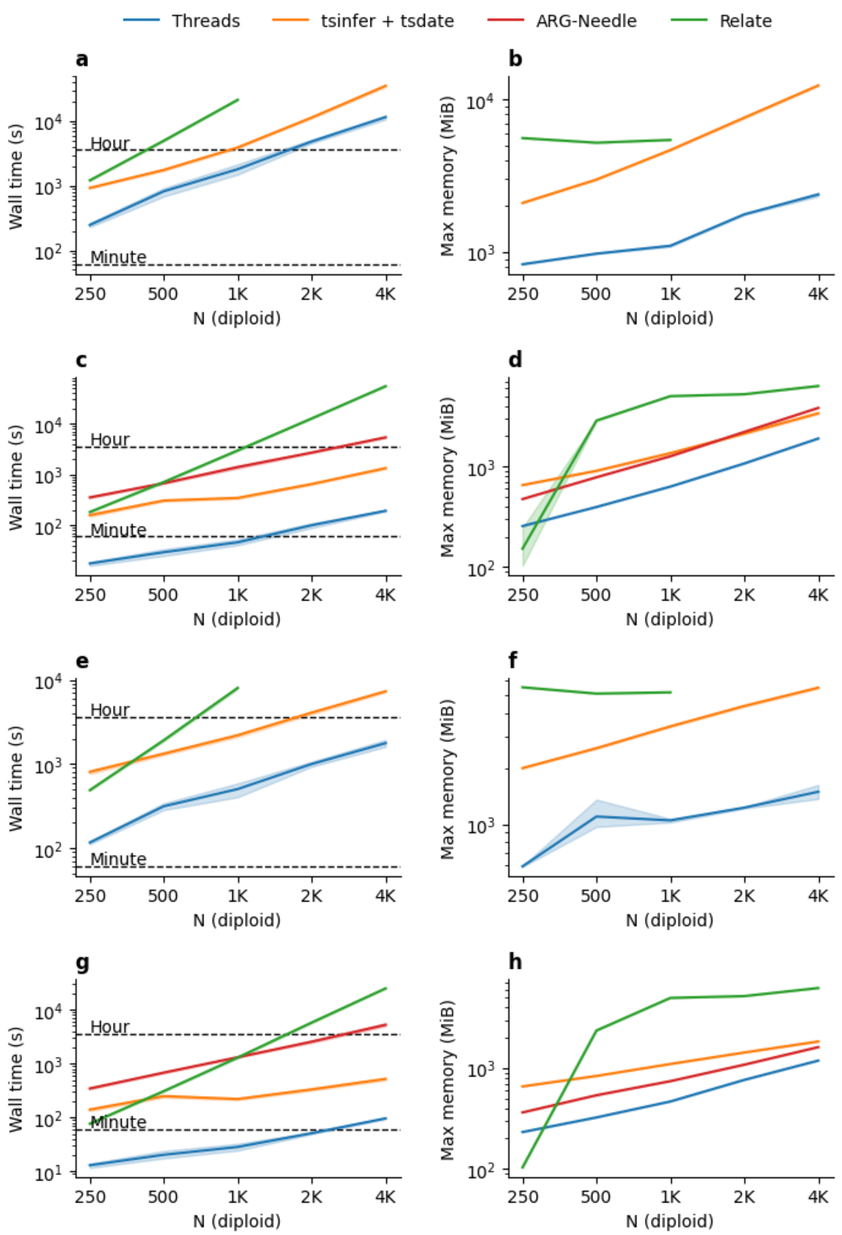


**Supplementary Figure 5. Computational performance using a single CPU**. We measure runtime, peak memory usage, and disk space required for final output. **a-b**. Simulated sequencing data under a European (CEU) demography. **c-d**. Simulated genotyping array data under a European (CEU) demography. **e-f.** Simulated sequencing data under a constant (Ne=10,000) demography. **g-h.** Simulated genotyping array data under a constant (Ne=10,000) demography. Shaded regions show 95% confidence intervals across 15 random seeds.

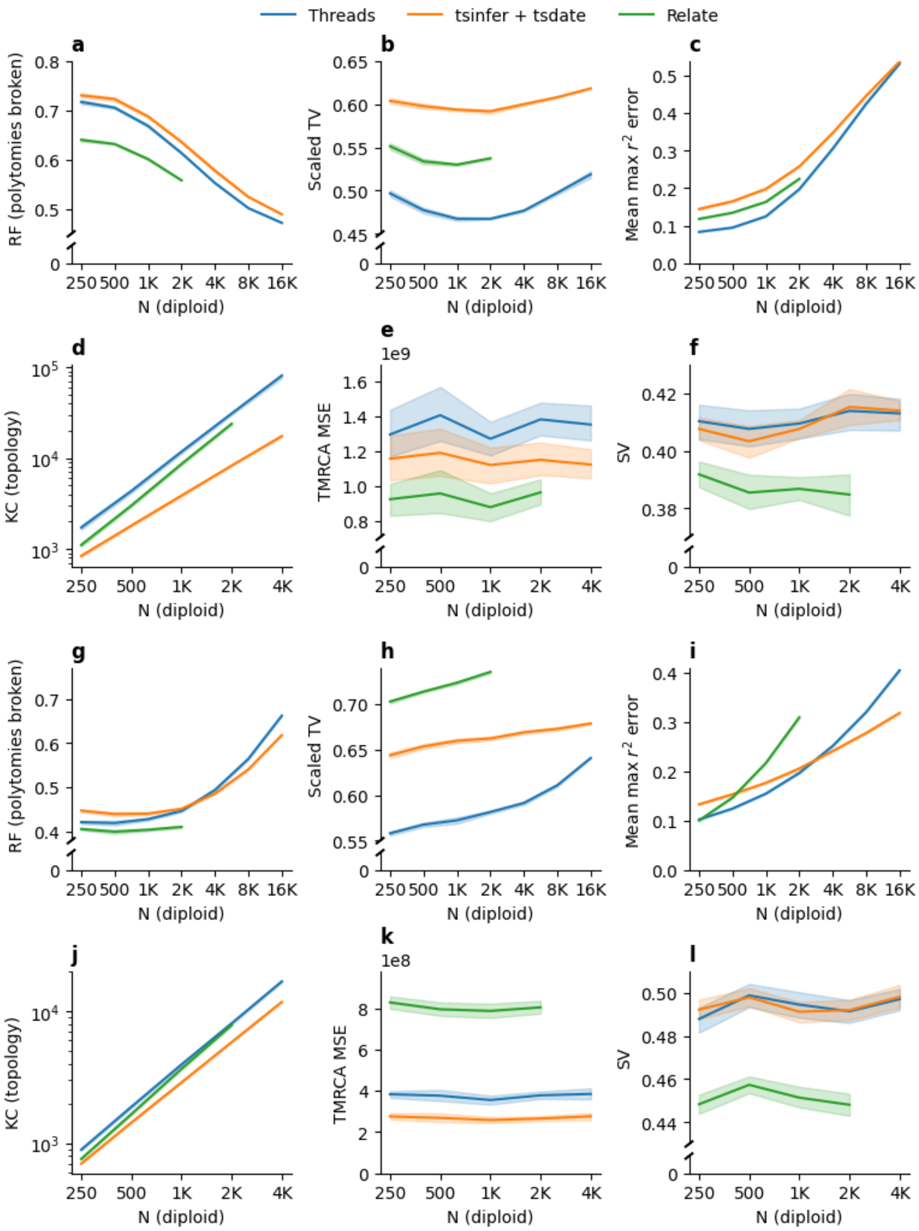


**Supplementary Figure 6. ARG inference accuracy on simulated sequencing data**. We evaluated six metrics, the Robinson-Foulds (RF), scaled total variation (TV), the mean max-$r^2$ error, the topological Kendall-Colijn (KC), the mean square error (MSE) and the split-size vector (SV). Shaded regions show 95% confidence intervals across 15 random seeds. **a-f**. Simulations under a European (CEU) demography. **g-l.** Simulations under a constant (Ne=10,000) demography.

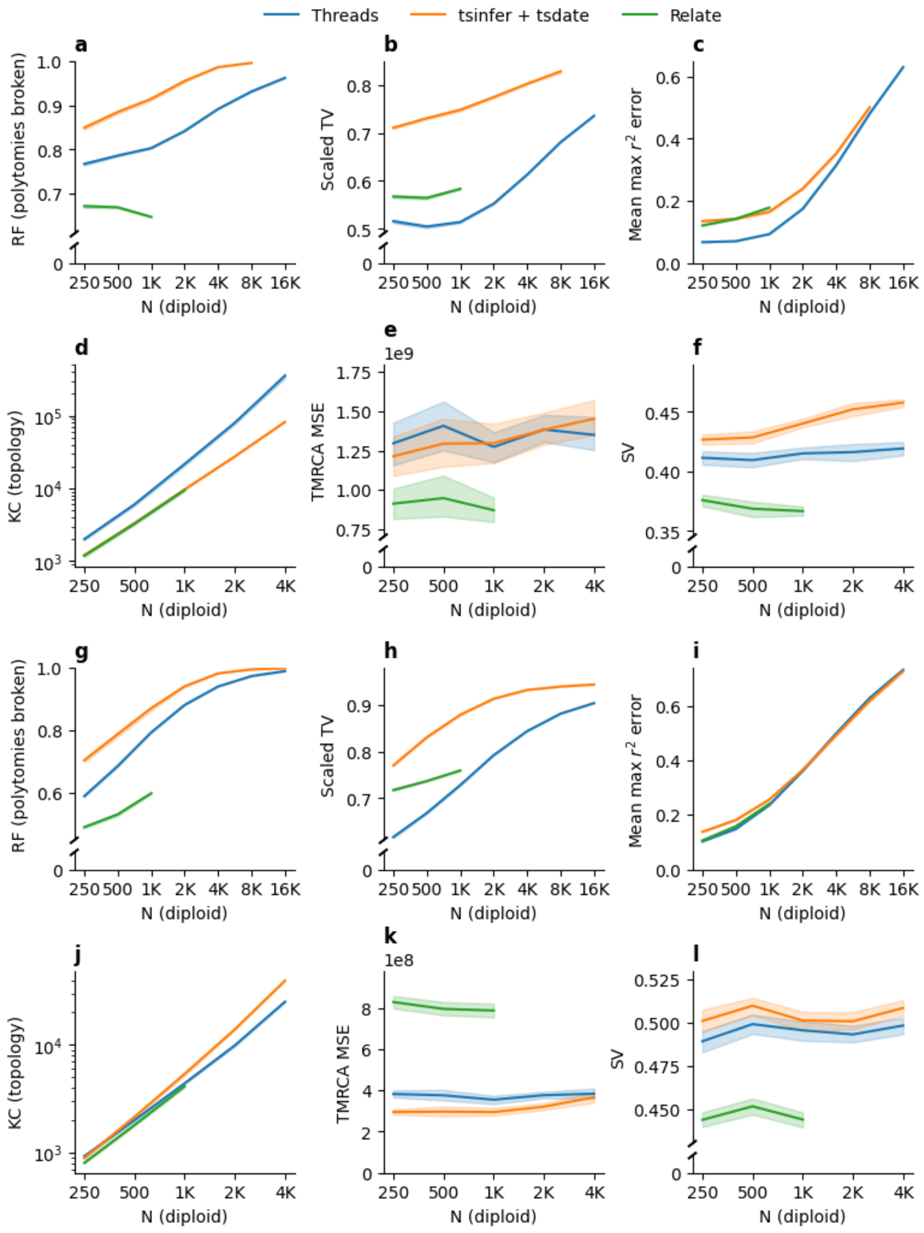


**Supplementary Figure 7. ARG inference accuracy on simulated sequencing data with a 0.1% error rate**. We evaluated six metrics, the Robinson-Foulds (RF), scaled total variation (TV), the mean max-r$^2$ error, the topological Kendall-Colijn (KC), the mean square error (MSE) and the split-size vector (SV). Shaded regions show 95% confidence intervals across 15 random seeds. **a-f**. Simulations under a European (CEU) demography. **g-l.** Simulations under a constant (Ne=10,000) demography.

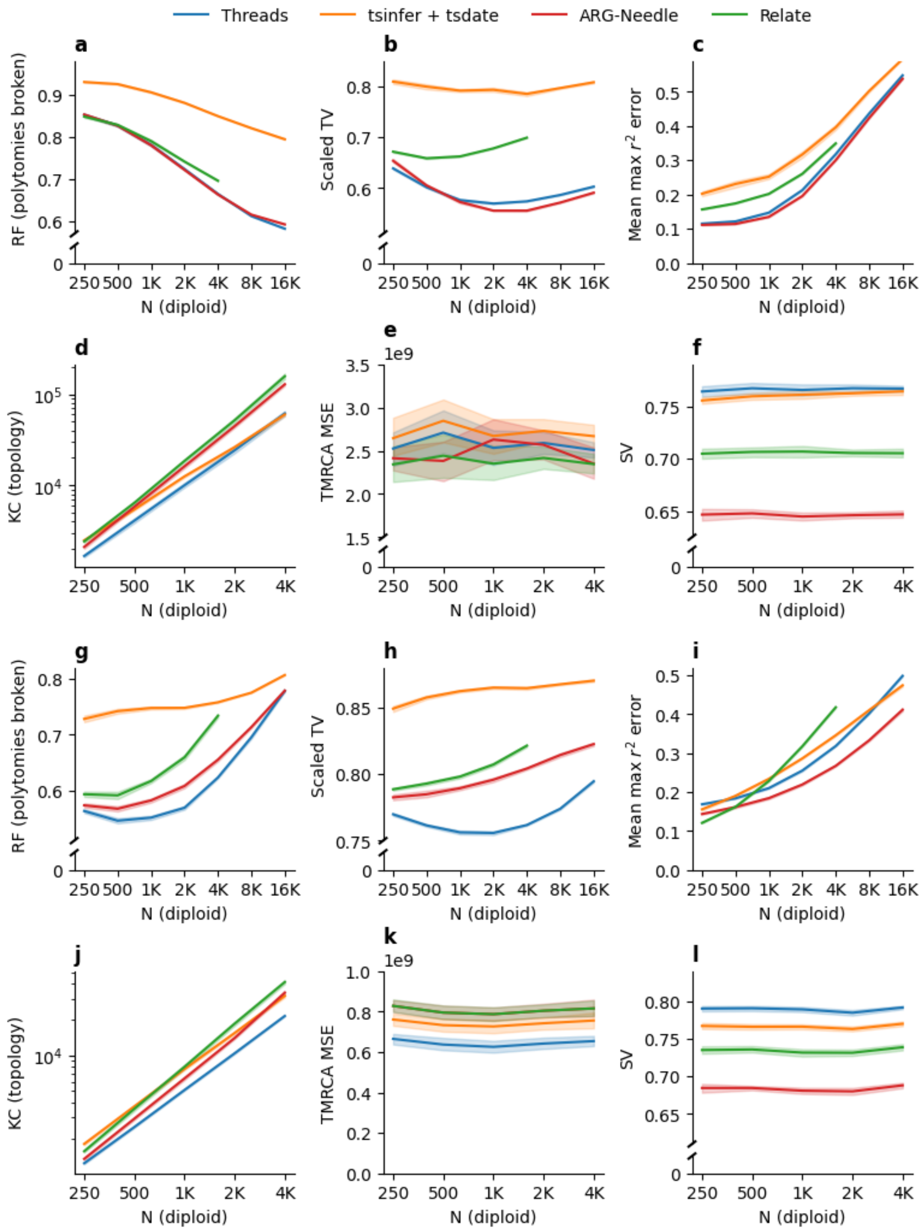


**Supplementary Figure 8. ARG inference accuracy on simulated genotyping array data**. We evaluated six metrics, the Robinson-Foulds (RF), scaled total variation (TV), the mean max-$r^2$ error, the topological Kendall-Colijn (KC), the mean square error (MSE) and the split-size vector (SV). Shaded regions show 95% confidence intervals across 15 random seeds. **a-f**. Simulations under a European (CEU) demography. **g-l.** Simulations under a constant (Ne=10,000) demography.

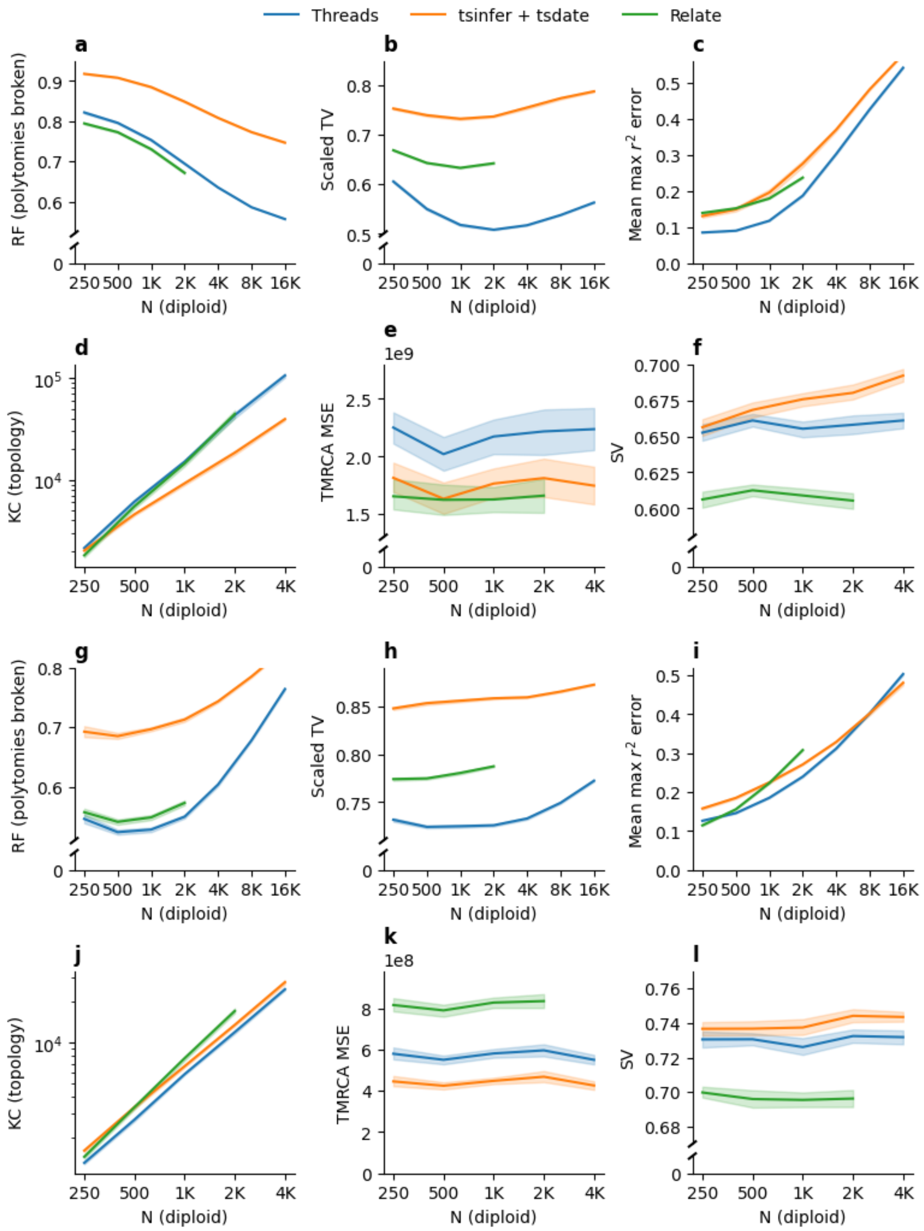


**Supplementary Figure 9. ARG inference accuracy on simulated sequencing data with a low mutation rate of $1.4\times10^{-9}$**. We evaluated six metrics, the Robinson-Foulds (RF), scaled total variation (TV), the mean max-$r^2$ error, the topological Kendall-Colijn (KC), the mean square error (MSE) and the split-size vector (SV). Shaded regions show bootstrap 95% confidence intervals across 15 random seeds. **a-f**. Simulations under a European (CEU) demography. **g-l.** Simulations under a constant (Ne=10,000) demography.

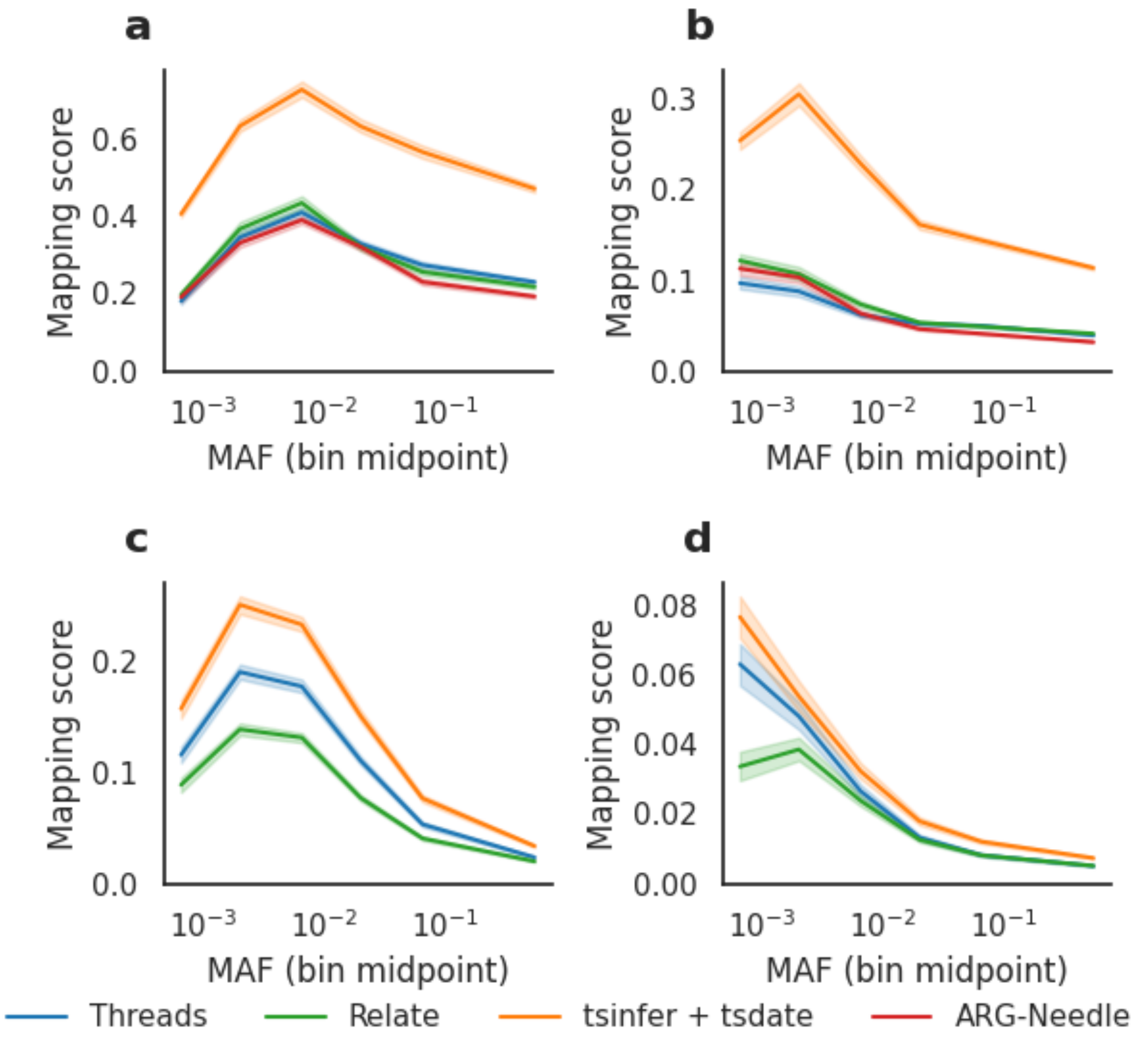


**Supplementary Figure 10. Mapping score in simulated array and sequencing data**. We simulated array (**a, b**) and sequencing data (**c, d**) for CEU (**a, c**) and constant N=10K (**b, d**) demographic models. The mapping score is computed as the number of mutations needed to heuristically map a genotype vector on an inferred marginal tree, divided by the derived allele count. In these experiments, we did not attempt to randomly resolve polytomies when running tsinfer+tsdate. Shaded regions show bootstrap 95% confidence intervals across 15 random seeds.

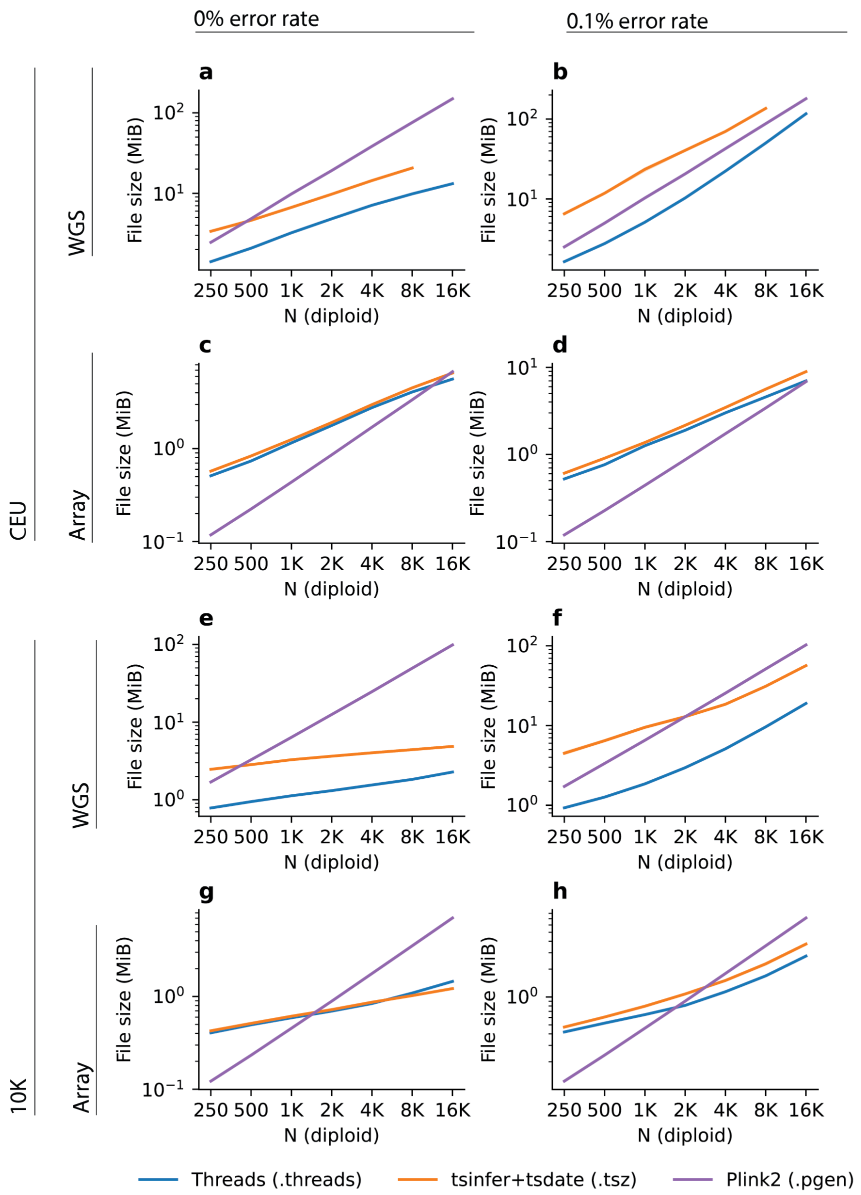


**Supplementary Figure 11. Disk space usage for data simulated data**. Simulations used a European demographic model and a constant-sized demographic model for sequencing and genotyping data, with and without errors. Threads and tsinfer+tsdate formats encode both genotype and genealogical data, PLINK2 stores only genotype data.

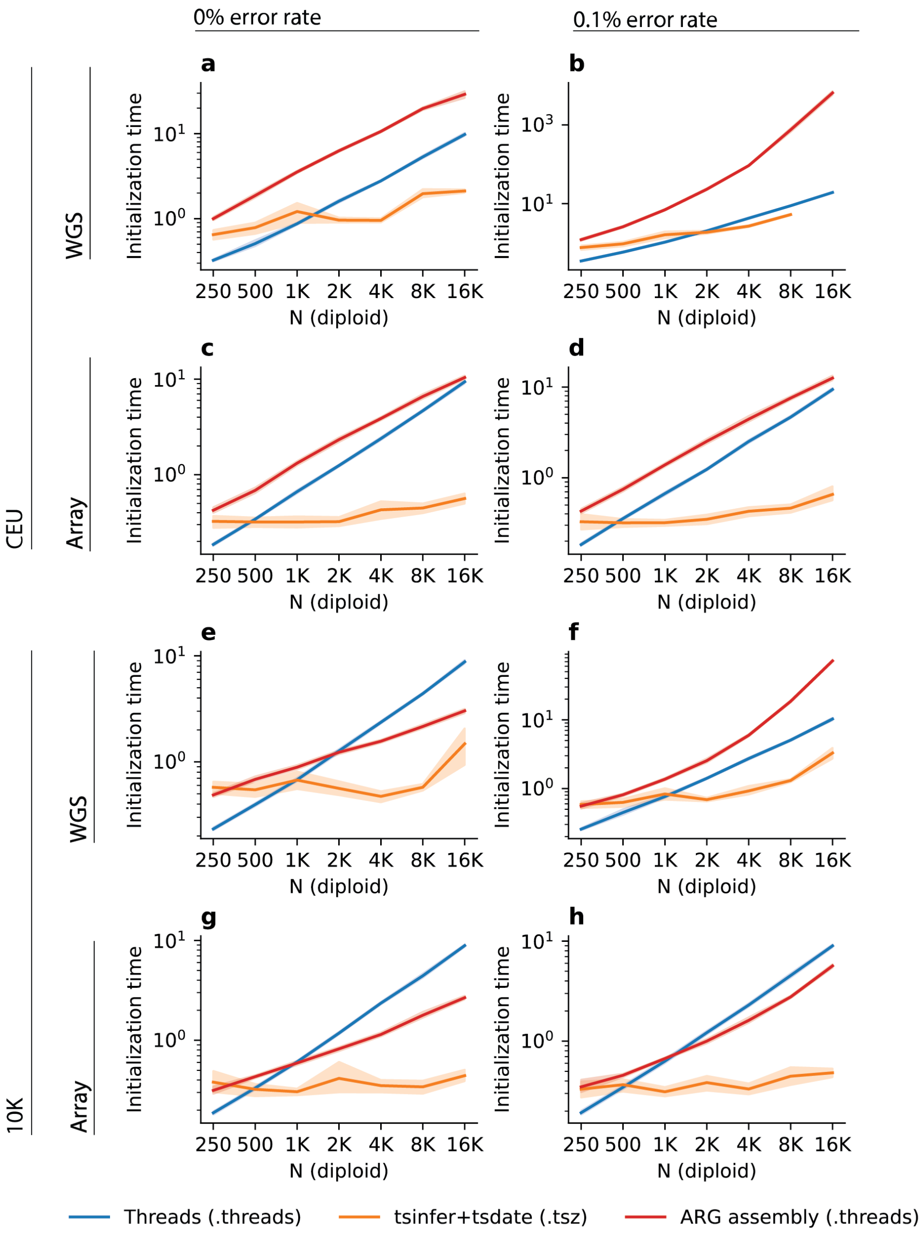


**Supplementary Figure 12. Time required to read different ARG-based data structures into memory from disk.** Results for data simulated under a European demographic model (CEU) and a constant-sized demographic model (10K) for simulated sequencing and genotyping array data, with and without errors. "Threads (.threads)" refers to building the graph shown in Supplementary Figure 1 from a .threads file containing Threads-compressed threading instructions. "tsinfer+tsdate (.tsz)" refers to loading an ARG inferred using tsinfer+tsdate and stored in tskit tree sequence format, decompressed using the tszip library. "ARG assembly (.threads)" refers to assembling an ARG using the arg-needle-lib library. Instead of loading an ARG in arg-needle-lib argn format, the ARG is built by executing the threading instructions contained in a .threads file.

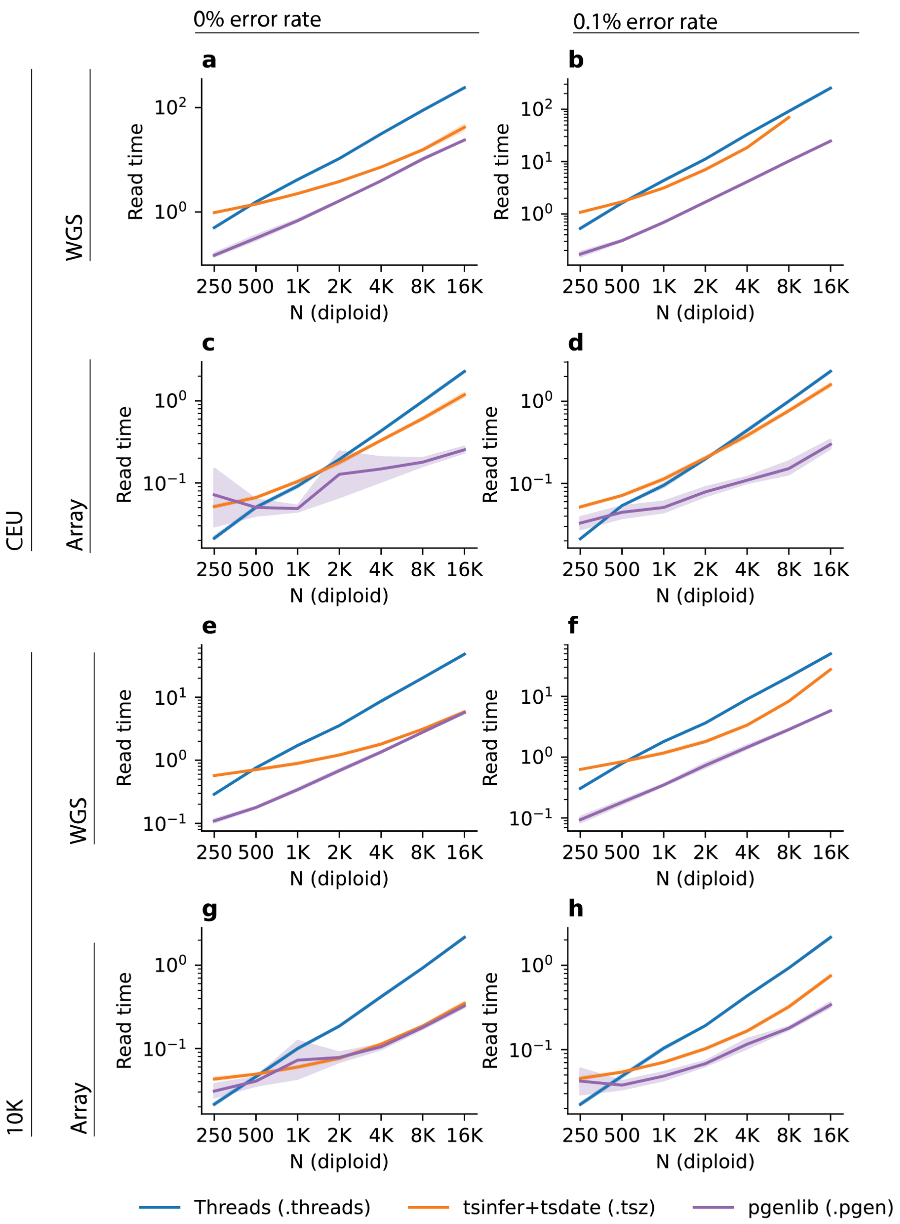


**Supplementary Figure 13. Time required to read a full genotype matrix into memory from initialized data structures.** We simulated individuals from a European demographic model (CEU) and a constant-sized demographic model (10K) for sequencing and SNP array data, with and without errors. We pre-loaded the .threads and .tsz files into memory as described in Supplementary Figure 18. “Threads (.threads)” refers to recovering the genotype matrix from the preloaded data structure described in Supplementary Figure 1. “tsinfer+tsdate (.tsz)” refers to extracting the genotype matrix from an ARG loaded in memory in tskit tree sequence format. “pgenlib” refers to loading the genotype matrix from disk, using the pgenlib library.

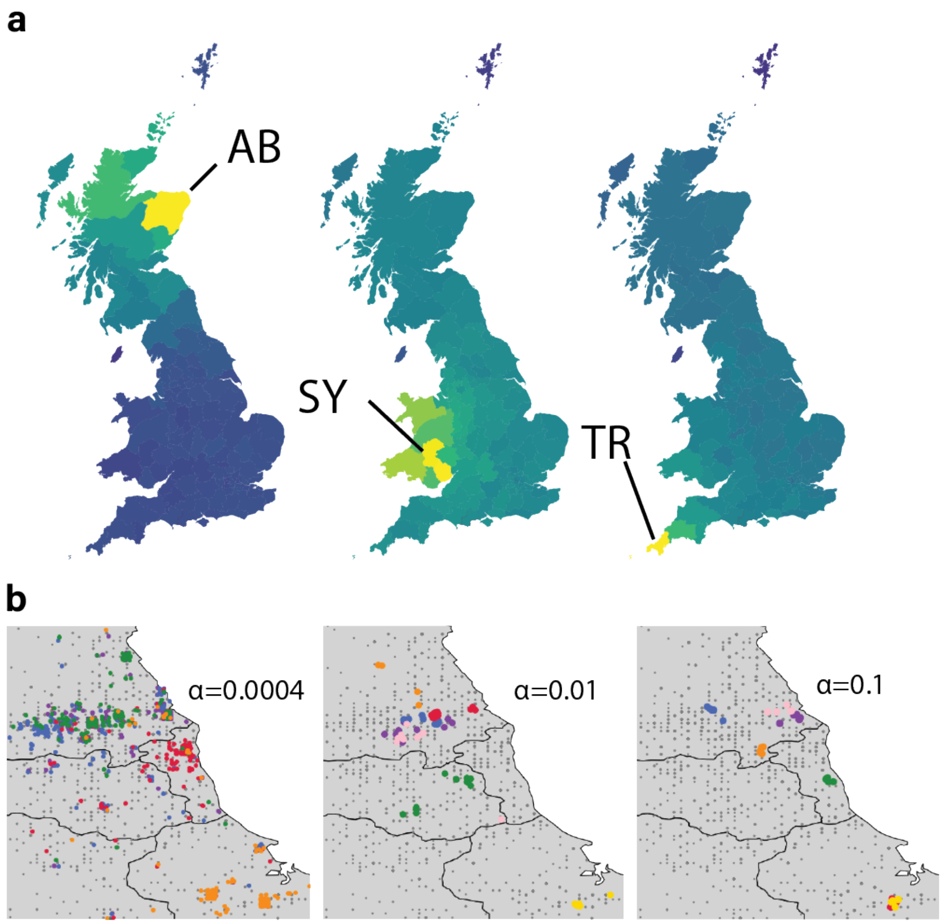


**Supplementary Figure 14. ARG-based fine-scale population structure analysis. a.** We explored the use of inferred ARGs to recover regional structure by ascertaining the geographic origin of genealogical closest cousins for 100 samples from each UK Biobank postcode. Brighter colors indicate a higher probability of finding genealogical closest cousins for samples from a focal postcode. We show the distributions for three example postcodes: Aberdeen (AB), Shrewsbury (SY), and Truro (TR). **b.** We performed clustering of samples from five postcodes in North-East England using the proportion of genome shared through IBD segments within 10 generations. The clustering was truncated for relatedness thresholds $\alpha = 0.0004, 0.1, 1.0$ (see Methods). Each panel shows results of extracting the largest 5, 7, and 7 clusters, respectively, with different clusters represented using different colors. Gray dots represent samples not belonging to any of the highlighted clusters.

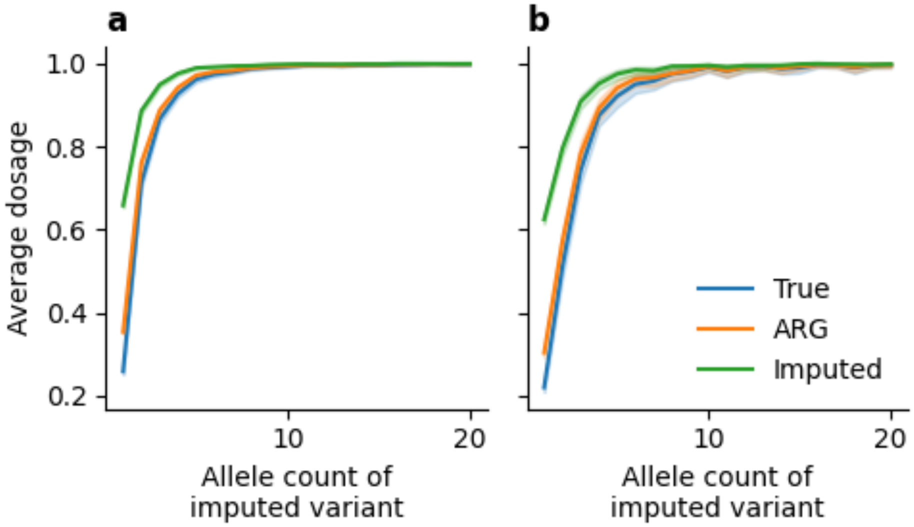


**Supplementary Figure 15. Inflation of imputation dosages in simulated data.** We simulated reference panels of size 100 (**a**) and 1,000 (**b**), assuming perfect inference of ARGs for ARG-based imputation and perfect inference of genealogical closest cousins for imputation. If any of the closest genealogical cousins of a target sequence being imputed carries a mutation, we estimate, through simulation, genotype dosages using both the ARG and imputation based on haplotype copying. We compare these with the true, simulated values, averaging across bins of minor allele count in the reference panel. The case "ARG" is different to the true genotype only when the target sequence coalesces directly into the lineage carrying the mutation being imputed, in which case we estimate the dosage based on the mutation being located uniformly at random along the lineage. For "Imputed", we average over the allele state of all genealogical closest cousins. Simulations were performed under a European (CEU) demographic model.

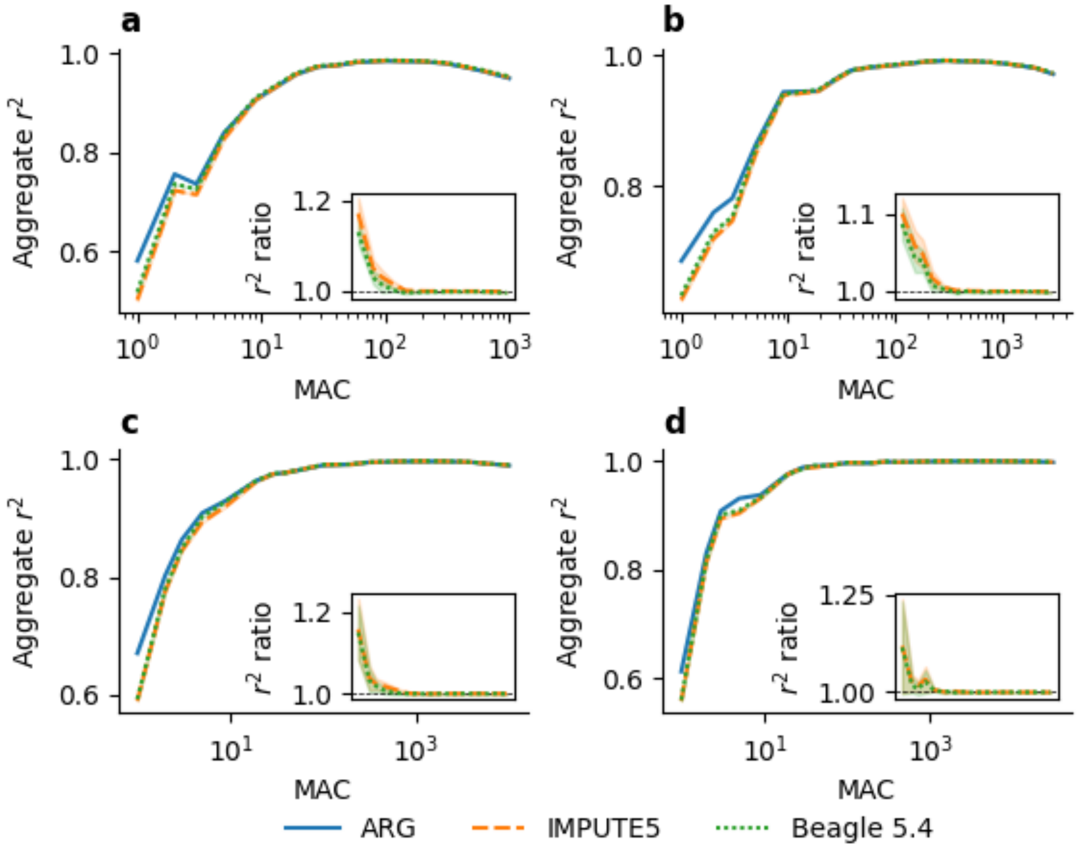


**Supplementary Figure 16. Imputation performance of Threads (ARG), IMPUTE5 and Beagle 5.4 using simulated reference panels**. We simulated panels of size 1,000 (**a**), 3,000 (**b**), 10,000 (**c**), and 30,000 (**d**), and measured accuracy using 10 held-out samples, stratifying the results by allele count in the panel. We measured Accuracy using aggregate $r^2$ of between imputed dosages and the true, simulated genotypes. Insets show the ratio between Threads and other methods.

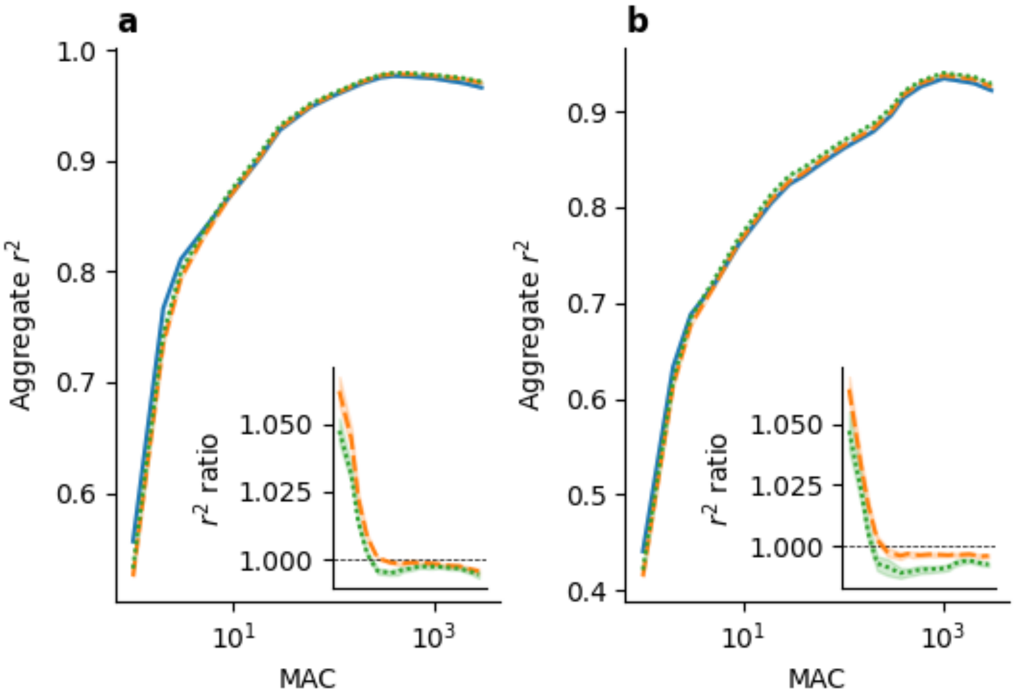


**Supplementary Figure 17. Imputation accuracy for 1000 Genomes Project samples**. We measured imputation accuracy using 7 samples of African (**a**) and European (**b**) ancestry. Inset plot shows the ratio between ARG-based imputation and IMPUTE5 or Beagle 5.4. **b.** Imputation quality for 10 held-out samples from 5 European subpopulations (CEU, FIN, GBR, IBS, TSI) from the 1000 Genomes Project, measured by aggregate $r^2$ stratified by minor allele count in the reference panel. The inset shows the ratio between Threads and the other two methods, using the same x axis range.

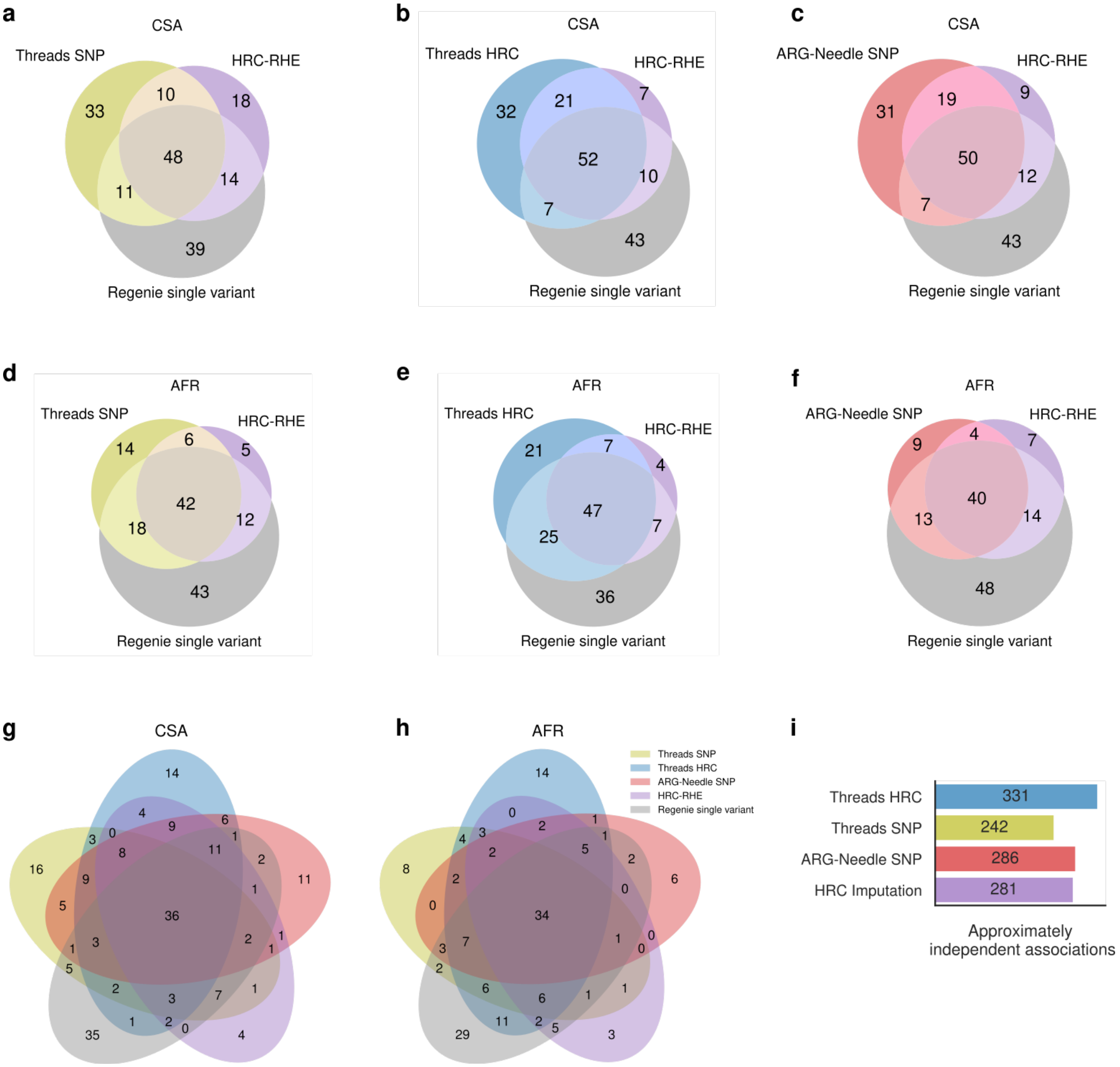


**Supplementary Figure 18. Additional details of association analyses. a-f.** Overlap in LD blocks containing gene-trait associations detected using HRC-RHE, Regenie, and ARG-RHE applied to ARGs inferred using either Threads (**a, b, d, e**) or ARG-Needle (**c, f**) in samples of Central/South Asian (CSA) ancestry (**a-c**) and African (AFR) ancestry (**d-f**). **g, h.** The total number of independently associated variants discovered using three association strategies for 52 quantitative traits in samples of **g.** CSA ancestry and **h.** AFR ancestry in the UK Biobank. **i.** Approximately independent single-variant signals in 52 quantitative traits detected through genealogy-wide association of ARGs inferred using Threads or ARG-Needle, as well as through a GWAS of HRC-imputed variants.

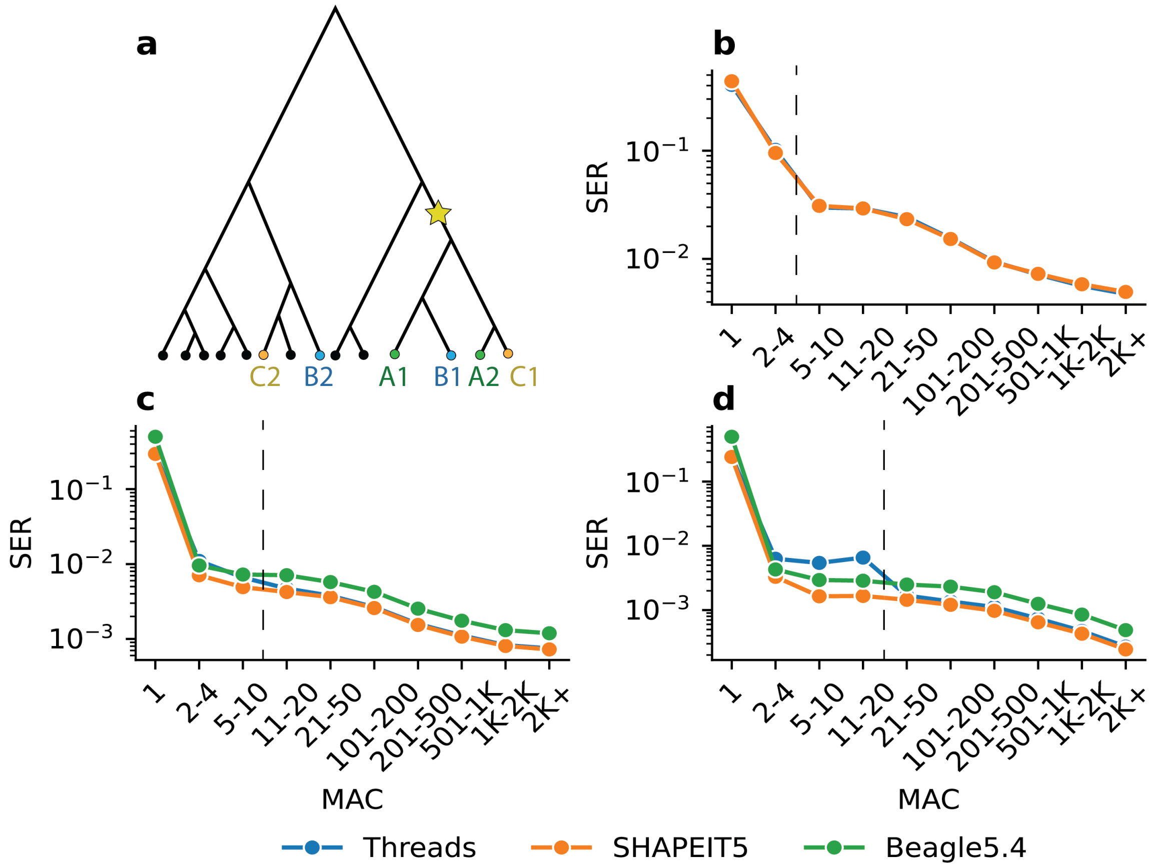


**Supplementary Figure 19. ARG-based phasing of rare-variants**. (**a**) A toy example of how ARGs may be used for phasing. Here, the mutation represented by a star symbol has three carriers, two heterozygous (B, C) and one homozygous (A). For each heterozygous carrier we evaluate which haplotype is more likely to be a carrier by observing how leaves in the marginal tree, which represent carriers, cluster together. (**b**) Phasing accuracy for a trio-phased sample (NA12878) in the 1000 Genomes Project for Threads and SHAPEIT5 measured using the switch error rate (SER) and stratified by MAC. (**c-d**) Phasing accuracy in simulations of size (**c**) N=3,000 and (**d**) N=10,000. Variants of allele frequency higher than 0.1% were phased using SHAPEIT5-common and used as a scaffold for both methods. Dashed vertical lines show the cut-off between rare and common-variant phasing methods.
